# Functionally Coherent Transcription Factor Target Networks Illuminate Control of Epithelial Remodelling

**DOI:** 10.1101/455709

**Authors:** Ian M. Overton, Andrew H. Sims, Jeremy A. Owen, Bret S. E. Heale, Matthew J. Ford, Alexander L. R. Lubbock, Erola Pairo-Castineira, Abdelkader Essafi

## Abstract

Cell identity is governed by gene expression, regulated by Transcription Factor (TF) binding at cis-regulatory modules. We developed the NetNC software to decode the relationship between TF binding and the regulation of cognate target genes in cell decision-making; demonstrated on nine datasets for the Snail and Twist TFs, and also modENCODE ‘HOT’ regions. Results illuminated conserved molecular networks controlling development and disease, with implications for precision medicine. Predicted ‘neutral’ TF binding accounted for the majority (50% to ≥80%) of candidate target genes from statistically significant peaks and HOT regions had high functional coherence. Expression of orthologous functional TF targets discriminated breast cancer molecular subtypes and predicted novel tumour biology. We identified new gene functions and network modules including crosstalk with notch signalling and regulation of chromatin organisation, evidencing networks that reshape Waddington’s landscape during epithelial remodelling. Predicted invasion role*s* were validated using a tractable cell model, supporting our computational approach.

## INTRODUCTION

Transcriptional regulatory factors (TFs) govern gene expression, which is a crucial determinant of phenotype. Mapping transcriptional regulatory networks is an attractive approach to understand the molecular mechanisms underpinning both normal biology and disease (Rhee et al., 2014; Shlyueva et al., 2014; Stampfel et al., 2015). TF action is controlled in multiple ways; including protein-protein interactions, DNA sequence affinity, 3D chromatin conformation, post-translational modifications and the processes required for TF delivery to the nucleus (Khoueiry et al., 2017; Rhee et al., 2014; Zabidi and Stark, 2016). A complex interplay of mechanisms influences TF specificity across different biological contexts and genome-scale assignment of TFs to individual genes is challenging (Khoueiry et al., 2017; Shlyueva et al., 2014; Wilczynski and Furlong, 2010).

TF binding sites may be determined using chromatin immunoprecipitation followed by sequencing (ChIP-seq) or microarray (ChIP-chip). These and related methods (e.g. ChIP-exo, DamID) have revealed a substantial proportion of statistically significant ‘neutral’ TF binding, that has apparently no effect on transcription from assigned target genes (Biggin, 2011; Li et al., 2008; Ozdemir et al., 2011; Shlyueva et al., 2014). Genomic regions that bind large numbers of TFs are termed Highly Occupied Target (HOT) regions (Roy et al., 2010), are enriched for disease SNPs and can function as developmental enhancers (Kvon et al., 2012; Li et al., 2015). However, a considerable proportion of individual TF binding events at HOT regions may have little effect on gene expression and association with chromatin accessibility suggests non-canonical regulatory function such as sequestration of TFs or in 3D genome organisation (Montavon et al., 2011; Moorman et al., 2006) as well as possible technical artefacts (Teytelman et al., 2013). Apparent neutral binding events may also have subtle functions; for example in combinatorial context-specific regulation or in buffering noise (Cannavò et al., 2016; Stampfel et al., 2015). Identification of *bona fide*, functional TF target genes remains a major obstacle in understanding the regulatory networks that control cell behaviour (Biggin, 2011; Brown and Celniker, 2015; Keung et al., 2014; Khoueiry et al., 2017; Stampfel et al., 2015).

Genes regulated by an individual TF typically have overlapping expression patterns and coherent biological function (Igual et al., 1996; Karczewski et al., 2014; MacArthur et al., 2009). Indeed, gene regulatory networks are organised in a hierarchical, modular structure and TFs frequently act upon multiple nodes of a given module (Hartwell et al., 1999; Hooper et al., 2007). Therefore, we hypothesised that functional TF targets have network properties that are different to neutrally bound sites. Network analysis can reveal biologically meaningful gene modules, including cross-talk between canonical pathways (Ideker et al., 2002; Jaeger et al., 2017; Vidal et al., 2011) and so may enable elimination of neutrally bound candidate TF targets derived from statistically significant ChIP-seq or ChIP-chip peaks. Network approaches afford significant advantages for handling biological complexity, enable genome-scale analysis of gene function (Greene et al., 2015; Hu et al., 2016), and are not restricted to predefined gene groupings used by standard functional annotation tools (e.g. GSEA, DAVID) (Huang et al., 2009; Ideker et al., 2002; Subramanian et al., 2005). Clustering is frequently applied to define biological modules (Enright et al., 2002; Wang et al., 2011). However, pre-defined modules may miss condition-specific features; for example, gene products may be absent in the biological condition(s) analysed but present in the base network. Hence, clusters derived from a whole-genome network may not accurately capture biological interactions that occur in a particular context. Context-specific interactions are common, for example the varied repertoire of biophysical interactions in different cell types or between cell states, such as in the stages of the cell cycle (Pawson and Nash, 2003). We developed a functional TF target discovery algorithm (NetNC) with capability for context-specific module discovery. The Snail and Twist TFs have important roles in Epithelial to Mesenchymal Transition (EMT), a multi-staged morphogenetic programme fundamental for normal embryonic development that contributes to tumour progression and fibrosis (Giampieri et al., 2009; Lim and Thiery, 2012; Nieto et al., 2016; Yu et al., 2013). We analysed Snail and Twist binding datasets, integrating with genetic screens and breast cancer transcriptomes to gain insight into epithelial remodelling in development and disease.

## RESULTS AND DISCUSSION

### A comprehensive *D. melanogaster* functional gene network (DroFN)

We developed a systems-wide map of *D. melanogaster* signalling and metabolism (DroFN; 11,432 nodes, 787,825 edges). DroFN performed well on time-separated blind test data (TEST-NET) compared with DroID (Yu et al., 2008) and GeneMania (Warde-Farley et al., 2010) (Table S1, Figure S1). The overlap between DroFN and the *Drosophila* proteome interaction map (DPiM (Guruharsha et al., 2011)) was highly significant (Fisher Exact Test *p*<10^−308^). DroFN and DPiM had 999 genes in common and 37.8% (2175/5747) of DroFN edges for these genes were also found in DPiM. The DroFN False Positive Rate (0.047) was close to the prior for functional interaction estimated from KEGG (0.044); thus, some DroFN false positives may represent *bona fide* interactions not annotated in KEGG. Overall, DroFN provides a useful genome-scale map of pathway comembership in *D. melanogaster*.

### Prediction of functional transcription factor binding

We developed the NetNC algorithm for genome-scale prediction of functional TF target genes (Figure 1). NetNC builds upon observations that TFs coordinately regulate multiple functionally related targets (Igual et al., 1996; Karczewski et al., 2014; MacArthur et al., 2009) and was calibrated for discovery of biologically coherent genes in noisy data. An edge-centric ‘Functional Target Identification’ (FTI) mode seeks to distinguish all biologically coherent gene pairs from functionally unrelated targets. The node-centric ‘Functional Binding Target’ (FBT) mode analyses degree-normalised Node Functional Coherence Scores (NFCS). Gold-standard data for NetNC development and validation took KEGG pathways to represent biologically coherent relationships, combined with ‘Synthetic Neutral Target Genes’ (SNTGs). NetNC was robust to variation in input dataset size and %SNTGs, outperforming HC-PIN (Wang et al., 2011) and MCL (Enright et al., 2002) (Figure 2, Table S2). In general, NetNC was more stringent, with lower False Positive Rate (FPR) and higher Matthews Correlation Coefficient (MCC) than HC-PIN. At the highest %SNTG, MCC values for NetNC-FTI were around 50% to 67% higher than those for HC-PIN and NetNC-FBT typically had lowest FPR. NetNC’s performance advantages were most prominent on data with ≥50% SNTGs (Figure 2) and all nine TF_ALL datasets were predicted to contain ≥50% neutrally bound targets (Figure 3). Given the performance advantage at ≥50% STNGs, NetNC appears the method of choice for analysis of genome-scale TF occupancy data. However, NetNC may be applied to multiple different tasks, including: identification of differentially expressed pathways and macromolecular complexes from functional genomics data; illuminating common biology among CRISPR screen hits in order to inform prioritisation of candidates for follow-up work (Shalem et al., 2014); and discovery of functional coherence in chromosome conformation capture data (4-C, 5-C), for example in enhancer regulatory relationships (Dostie et al., 2006; Simonis et al., 2006).

**Figure 1.**
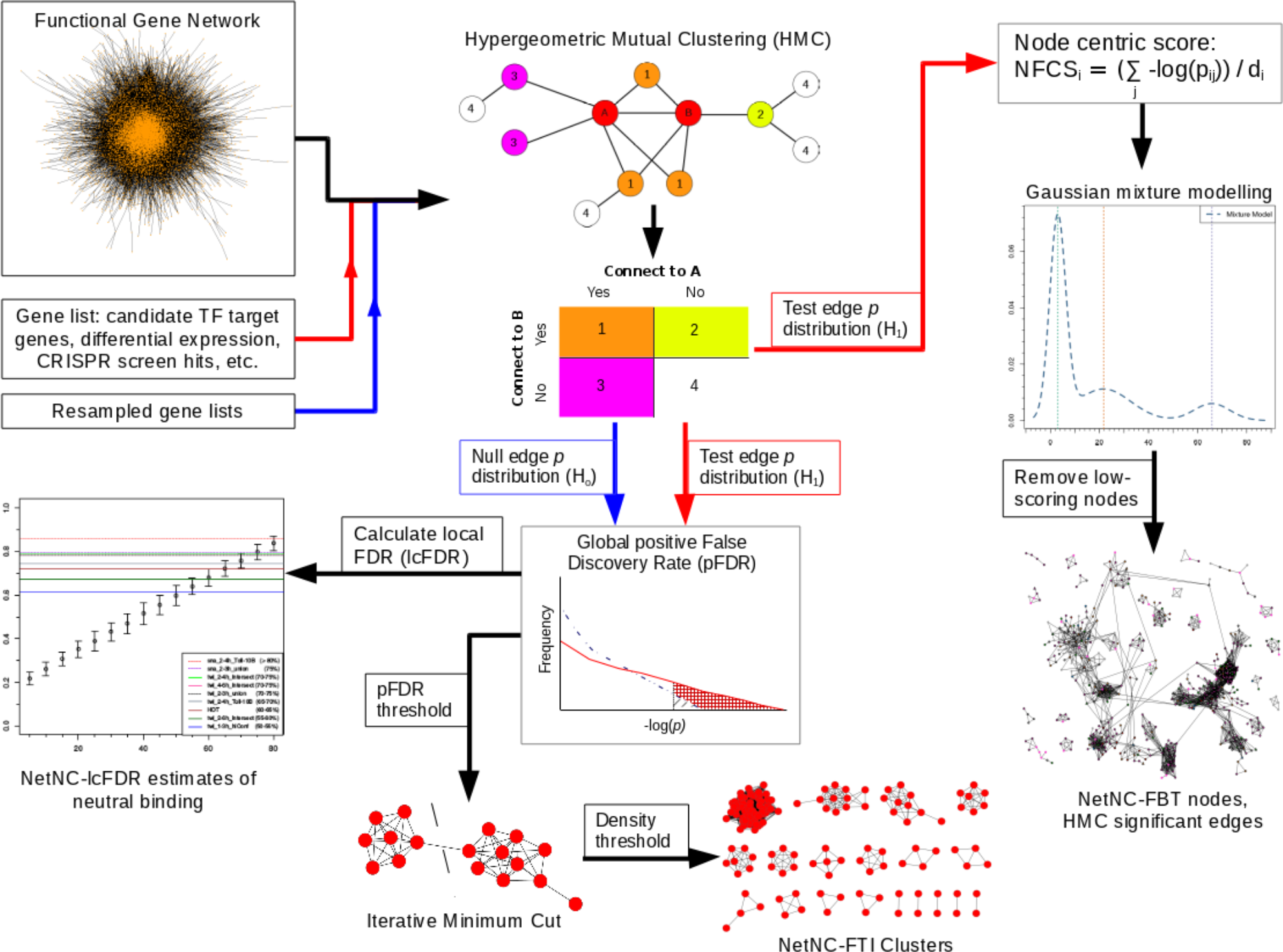
Overview of the NetNC algorithm. NetNC input data may be candidate TF target genes and a functional gene network (top, left). However, NetNC may be applied to any gene list and reference network. Hypergeometric Mutual Clustering (HMC) p-values are calculated for candidate TF target genes (top, middle); the node numbers and colours in the HMC graph correspond to those in the contingency table. HMC p-values are then employed in either i) a node-centric analysis mode (NetNC-FBT, right top) or ii) an edge-centric mode (NetNC-FTI) using global False Discovery Rate (pFDR, middle) followed by iterative minimum cut (bottom). NetNC can estimate local FDR (lcFDR) to predict the proportion of neutrally bound candidate target genes (left). NetNC can produce pathway-like clusters and also biologically coherent node lists for which edges may be taken using a standard FDR or Family Wise Error Rate (FWER) threshold on the HMC p-values (right).

**Figure 2.**
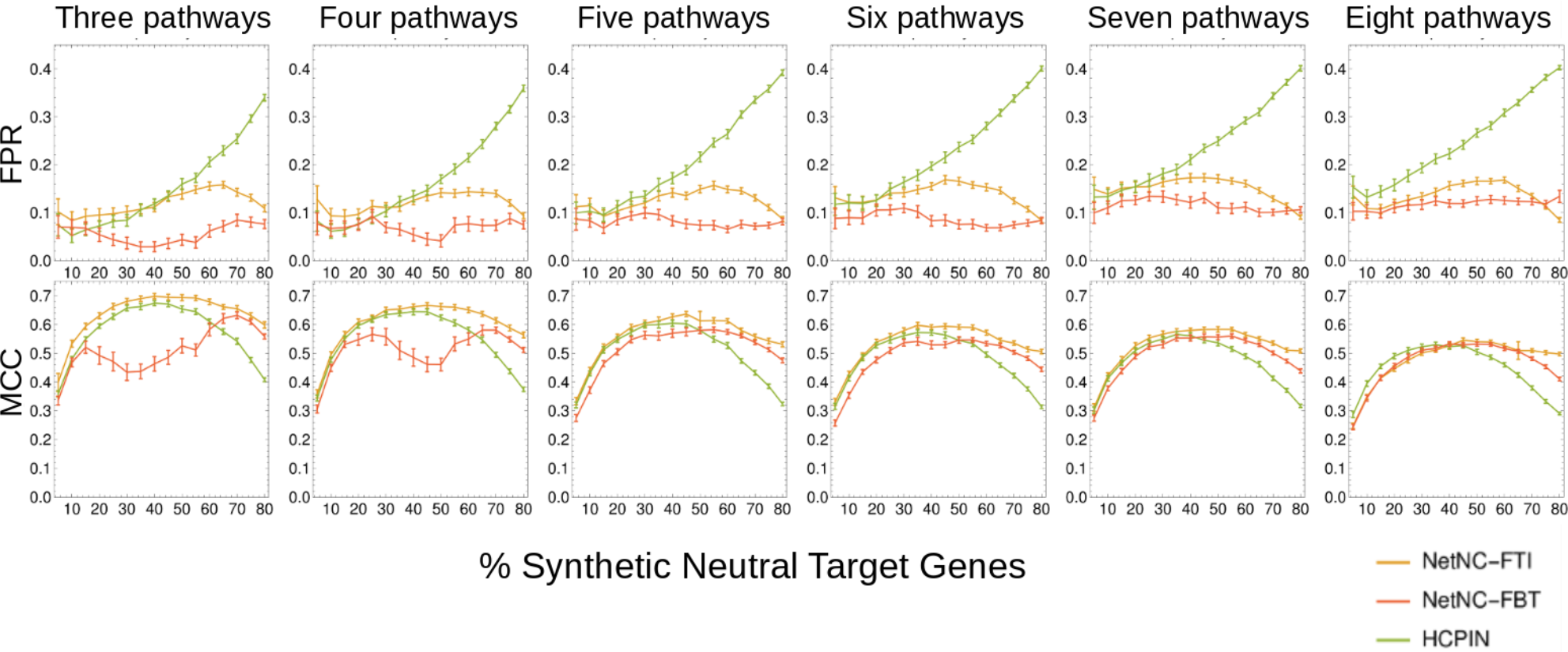
Evaluation of NetNC and HC-PIN on blind test data. Performance values reflect discrimination of KEGG pathway nodes from Synthetic Neutral Target Genes (STNGs), shown for NetNC-FTI (orange), NetNC-FBT (red) and HC-PIN (green). False Positive Rate (FPR, top row) and Matthews Correlation Coefficient (MCC, bottom row) values are given. Data shown represents analysis of TEST-CL_ALL, which included subsets of three to eight pathways, shown in columns, and sixteen %STNG values were analysed (5% to 80%, x-axis). NetNC performed best on the data examined with typically lower FPR and higher MCC values. Error bars reflect 95% confidence intervals calculated from quantiles of SNTG resamples. Also see Table S2.

**Figure 3.**
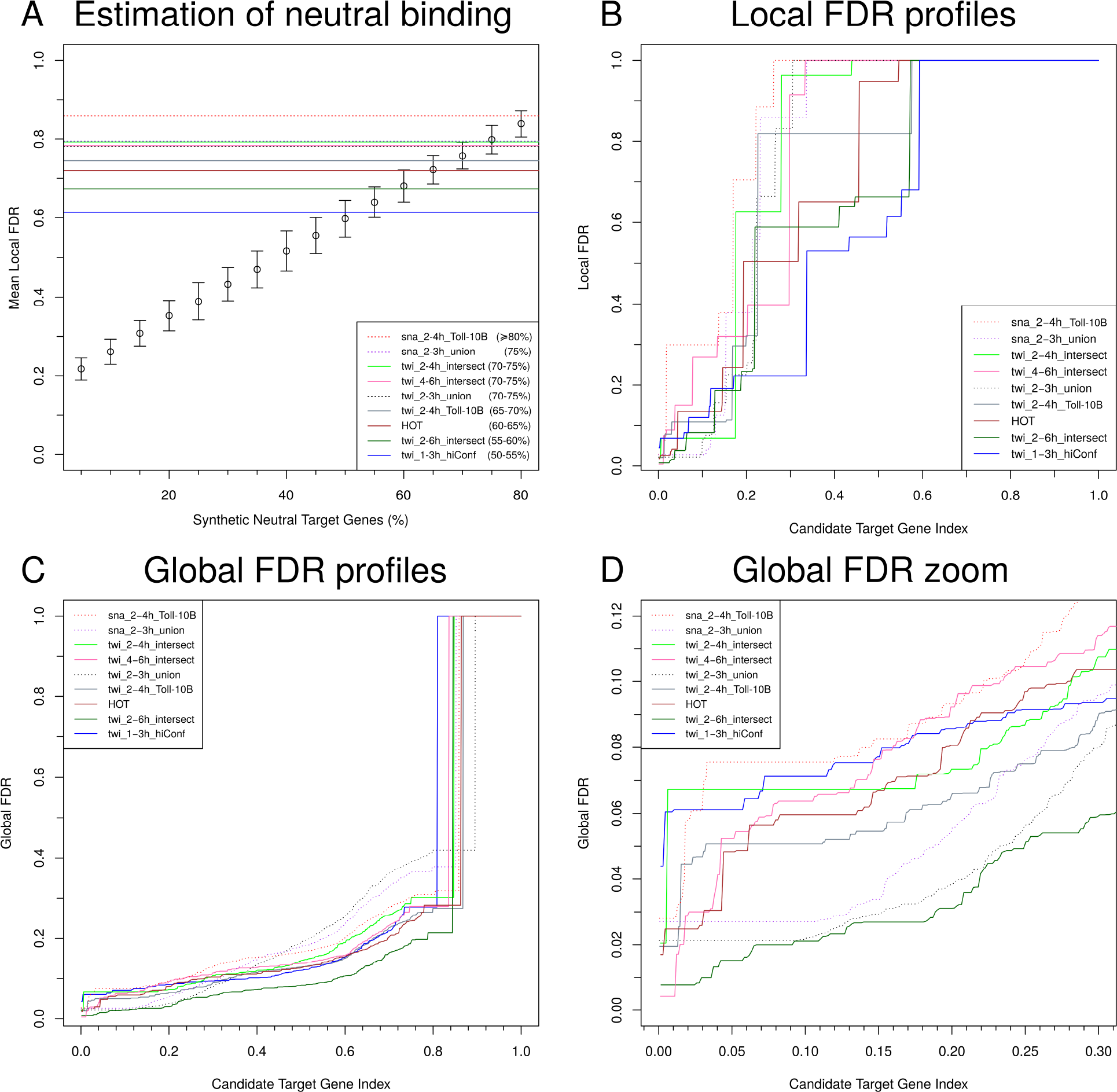
Neutral transcription factor binding and false discovery rate (FDR) profiles. Panel A: Estimation of total neutral binding. Black circles show NetNC mean lcFDR values for the TEST-CL_8PW data, ranging from 5% to 80% SNTGs; error bars represent 95% CI calculated from quantiles of the SNTG resamples. Coloured horizontal lines show mean NetNC-lcFDR values for the TF_ALL datasets. Comparison of the known TEST-CL_8PW %SNTG values with estimated total neutral binding values from mean NetNC-lcFDR showed systematic overestimation of neutral binding. Cross-referencing mean NetNC-lcFDR values for TF_ALL with those for TEST-CL_8PW gave estimates of neutral binding between 50% and ≥80% (see panel key). **Panels B, C and D.** Line type and colour indicates dataset identity (see key). Candidate target gene index values were normalised from zero to one in order to enable comparison across the TF_All datasets.

NetNC identifies coherent genes according to the functional gene network (FGN) structure. Statistical evaluation of network coherence, including FDR estimation, is available within NetNC for numerical thresholding. Therefore, NetNC is generally applicable to high-dimensional data including analysis of single-subject datasets, which is an important emerging area for precision medicine (Vitali et al., 2017). Application of statistical and graph theoretic methods for quantitative evaluation of relationships between genes (nodes) in NetNC offers an alternative to the classical emphasis on individual genes in studying the relationship between genotype and phenotype. Gold standard datasets and DroFN are available from the Biostudies database, NetNC is available from https://github.com/IanOverton/NetNC.git.

### Estimating neutral binding for EMT transcription factors and Highly Occupied Target (HOT) regions

We predicted functional target genes for the Snail and Twist TFs for developmental stages around gastrulation in *D. melanogaster*; analysing Chromatin ImmunoPrecipitation (ChIP) microarray (ChIP-chip) or sequencing (ChIP-seq) data for overlapping time periods in early embryogenesis produced by four different laboratories and also modENCODE Highly Occupied Target (HOT) regions (MacArthur et al., 2009; Ozdemir et al., 2011; Roy et al., 2010; Sandmann et al., 2007; Zeitlinger et al., 2007). Nine datasets were studied (TF_ALL, Table 1; please see Methods for details). NetNC-lcFDR predicted neutral binding proportion (PNBP) across TF_ALL ranged from 50% to ≥80% (Figure 3A, Table 1). Reassuringly, the most stringent peak calling approach (twi_1-3h_hiConf (Ozdemir et al., 2011) had the lowest (NetNC-lcFDR) or second lowest (NetNC-FTI) PNBP. Despite having lowest overall PNBP, twi_1-3h_hiConf also had the smallest proportion of genes passing pFDR<0.05 (Figure 3D). Targets bound during two consecutive developmental time periods (twi_2-6h_intersect (Sandmann et al., 2007)) also ranked highly, and so did HOT regions despite depletion for known TF motifs (Boyle et al., 2014; Chen et al., 2014; Kvon et al., 2012) (Figure 3A, Table 1). Indeed, twi_2-6h_intersect had significantly greater PNBP (binomial *p<*4.0×10^−15^) than datasets from the same study that represented a single time period (twi_2-4h_intersect, twi_4-6h_intersect) (Sandmann et al., 2007) (Figure 3). Therefore, predicted functional binding was enriched for regions occupied at >1 time period or by multiple TFs and results supported the emerging picture of widespread combinatorial control involving TF-TF interactions, cooperativity and TF redundancy (Jolma et al., 2015; Khoueiry et al., 2017; Long et al., 2016; Spitz and Furlong, 2012; Stampfel et al., 2015).

**Table 1.** Predicted Functional binding for Snail, Twist and HOT candidate target genes. Results for NetNC are given based on ‘Functional Target Identification’ (NetNC-FTI) and mean local FDR (NetNC-lcFDR) calibrated against datasets with a known proportion of resampled Synthetic Neutral Target Genes (SNTG) described in Methods section 4.2.3. The above datasets correspond to the following developmental stages: 2-4h stages 4-9 (except ‘2-4h_intersect datasets which were stages 5-7 (Sandmann et al., 2007)); 2-3h stages 4-6; 1-3h stages 2-6; 4-6h stages 8-9 (Sandmann et al., 2007); 0-12h stages 1-15; gastrulation occurs at stage 6 (Campos-Ortega and Hartenstein, 1997). Also see Figure S2.

| Dataset |  |  | Predicted functional targets |  |
| --- | --- | --- | --- | --- |
| Name | Developmental time period(s) | Candidate target genes* | NetNC-FTI | NetNC-lcFDR |
| twi_1-3h_hiConf | 1-3h | 664 | 202 (30%) | 45-50% |
| twi_2-6h_intersect | 2-4h and 4-6h | 615 | 241 (39%) | 40-45% |
| twi_2-4h_intersect | 2-4h only (not 4-6h) | 801 | 182 (23%) | 25-30% |
| twi_4-6h_intersect | 4-6h only (not 2-4h) | 818 | 126 (15%) | 25-30% |
| HOT | 0-12h<sup>+</sup> | 677 | 174 (26%) | 35-40% |
| twi_2-3h_union | 2-3h | 1848 | 424 (23%) | 25-30% |
| sna_2-3h_union | 2-3h | 1158 | 226 (20%) | 25% |
| twi_2-4h_Toll<sup>10b</sup> | 2-4h | 1238 | 279 (23%) | 30-35% |
| sna_2-4h_Toll<sup>10b</sup> | 2-4h | 1488 | 211 (14%) | ≤20% |

\*number of candidate targets that mapped to DroFN nodes.
<sup>+</sup>multiple time periods and 41 different TFs.

PNBP was similar for sites derived from either the union or intersection of two Twist antibodies, although the NetNC-FTI method found a higher number of functional targets for the intersection of antibodies (30.5% (116/334) vs 23% (424/1848)). Hits identified by multiple antibodies may be technically more robust due to reduced off-target binding (Sandmann et al., 2007). However, taking the union of candidate binding sites could eliminate false negatives arising from epitope steric occlusion due to protein interactions. Similar PNBP for either the intersection or the union of Twist antibodies suggests that, despite expected higher technical specificity, the intersection of candidate targets may not enrich for functional binding sites at the 1% peak-calling FDR threshold applied in (MacArthur et al., 2009; Sandmann et al., 2007). Fewer false negatives implies recovery of numerically more functional TF targets, likely producing denser clusters in DroFN, which could facilitate functional target detection by NetNC. Indeed, datasets representing the union of two antibodies ranked highly in terms of both the total number and proportion of genes recovered at lcFDR<0.05 or pFDR<0.05 (Figure 3). Even datasets with high PNBP had candidate target genes that passed stringent NetNC FDR thresholds; for example, sna_2-3h_union, twi_2-3h_union respectively had the highest and second-highest proportion of candidate targets at lcFDR<0.05 (Figure 3B). We found no evidence for benefit in using RNA polymerase binding data to guide allocation of peaks to candidate target genes (datasets sna_2-3h_union, twi_2-3h_union). The twi_2-4h_Toll^10b^, sna_2-4h_Toll^10b^ datasets had a relatively low peak threshold (two-fold enrichment), which may contribute to the high PNBP for sna_2-4h_Toll^10b^. We note that PNBP values might systematically overestimate neutral binding because some functional targets could be missed; for example due to errors in assigning enhancer binding to target genes and in *bona fide* regulation of genes that have few DroFN edges with other candidate targets. Additionally, calibration of lcFDR values against KEGG synthetic data might influence neutral binding estimates, due to potential differences in network properties between TF targets and KEGG pathways. Reasurringly, NetNC-lcFDR and NetNC-FTI neutral binding estimates showed good agreement (Table 1, Figure S2).

ChIP peak intensity putatively correlates with functional binding, although some weak binding sites are functional (Biggin, 2011; Chen et al., 2013). NetNC NFCS values and ChIP peak enrichment scores were significantly correlated in 6/8 datasets (*q*<0.05, HOT regions not analysed). Datasets with no significant correlation (twi_1-3h_hiConf, twi_2-6h_intersect) derived from protocols that enrich for functional targets and had lowest PNBP values (Figure 3A). Indeed, the median peak score for twi_2-6h_intersect was significantly higher than datasets taken from a single time period in same study (twi_2-4h_intersect, *q*<5.0×10^−56^; twi_4-6h_intersect, *q<*4.8×10^−58^). Predicted targets were enriched for human orthologues, for example 72% (453/628) of twi_2-3h_union NetNC-FBT functional targets had human orthology, significantly higher than the full twi_2-3h_union dataset (50%, 616/1220; *p*<3×10^−28^ binomial test). Enrichment for evolutionary conservation in Snail and Twist functional targets aligns with their regulation of fundamental developmental processes.

### Genome-scale functional transcription factor target networks

NetNC results offer a global representation of the mechanisms by which Snail and Twist exert tissue-specific regulation in early *D. melanogaster* embryogenesis (Figure 4, Figure S3, Supplemental File 1). Networks were annotated using GO and FlyBase (Ashburner et al., 2000; Gramates et al., 2017; Huang et al., 2009; Maere et al., 2005), revealing eleven biological groupings common to ≥4/9 TF_ALL datasets (Table S3). NetNC results were robust to subsampled input and, as expected, higher variance was found at lower subsampling rates (Table S4).

**Figure 4.**
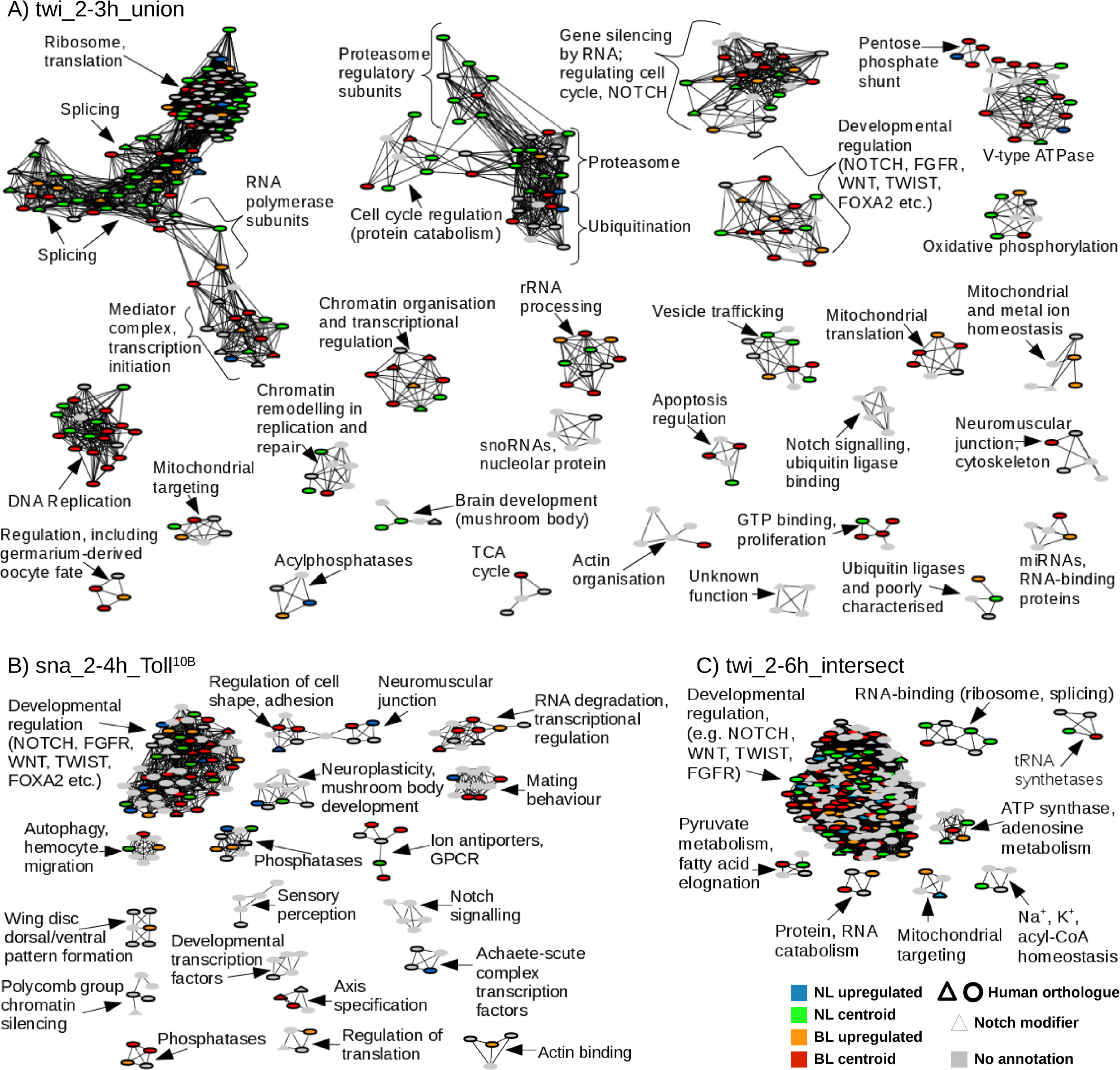
NetNC-FTI functional target networks for Snail and Twist. The key (bottom right) indicates annotations for human orthology (bold node border) and *Notch* screen hits (triangular nodes). Orthologues assigned basal-like (BL) or normal-like centroids (NL) are shown in red, green respectively; otherwise, node colour indicates differential expression NL (blue) vs BL (orange) subtypes (*q*<0.05) or no annotation (grey). Also see Figure S3, Tables S3 to S6 and Supplemental File 1. **Panel A: twi_2-3h_union.** Predicted functional targets cover several areas of fundamental biochemistry including splicing, DNA replication, energy metabolism, translation and chromatin organisation. Regulation of multiple conserved processes by Twist is consistent with the extensive cell changes required during mesoderm development. Clusters annotated predominantly to either NL or BL subtypes include mitochondrial translation (BL) and the proteasome (NL). These results predict novel Twist functions, for example in regulation of mushroom body neuroblast proliferation factors. **Panel B: sna_2-4h_Toll^10b^.** Multiple clusters of transcription factors were identified, aligning with Snail’s role as a master transcriptional regulator. Orthologues in the clusters ‘RNA degradation, transcriptional regulation’; ‘axis specification’ and ‘phosphatases’ were annotated exclusively to the basal-like subtype. **Panel C: twi_2-6h_intersect.** Most predicted functional targets belonged to the ‘developmental regulation’ cluster; dominance of regulatory factors might arise from the criterion for continuous binding across two developmental time windows. The developmental regulation cluster contained *mrr*, the orthologue of *IRX4*, which was upregulated in BL tumours.

NetNC analysis found Snail and Twist regulation of multiple core cell processes that govern the global composition of the transcriptome and proteome (Figure 4, Figure S3). These processes included transcription, chromatin organisation, RNA splicing, translation and protein turnover. A ‘Developmental Regulation Cluster’ (DRC) was identified in every TF_ALL dataset and contained members of multiple key conserved morphogenetic pathways, including notch, wnt and fibroblast growth factor (FGF). *Notch* signalling modifiers from public data (Guruharsha et al., 2012) overlapped significantly with NetNC-FTI results for each TF_ALL dataset (*q* <0.05), including the DRC, chromatin organisation and mediator complex clusters (Figure 4, Figure S3). *Notch* was an important control node across TF_ALL, with highest betweenness centrality in the DRC for three datasets and ranked among the top ten DRC genes for 8/9 datasets. Activation of *Notch* can result in diverse, context-specific transcriptional outputs and the mechanisms regulating this pleiotropy are not well understood (Bray, 2016; Guruharsha et al., 2012; Nowell and Radtke, 2017; Ntziachristos et al., 2014). Our results provided functional context for many *Notch* modifiers and proposed signalling crosstalk mechanisms in cell fate decisions driven by Snail and Twist, where regulation of modifiers may control the consequences of *Notch* activation. Crosstalk between *Notch* and *twist* or *snail* was previously shown in multiple systems, for example in adult myogenic progenitors (Bernard et al., 2010) and hypoxia-induced EMT (Sahlgren et al., 2008). Consistent with previous studies (Guruharsha et al., 2012; Ntziachristos et al., 2014), our results predict targeting of *Notch* transcriptional regulators, trafficking proteins, post-translational modifiers, receptor recycling and regulation of pathways that may attenuate or modify *Notch* signalling. Clusters where multiple modifiers were identified may represent cell meso-scale units important for *Notch* in the context of mesoderm development and EMT. For example, the mediator complex and transcription initiation subcluster for twi_2-3h_union (Figure 4) had thirteen nodes, of which five were *Notch* modifiers including orthologues of *MED7, MED8, MED31*. *Wingless* also frequently had high betweenness, ranking within the top ten DRC genes in six TF_ALL datasets and was twice top-ranked. Therefore *Notch* and *wingless* were key control points regulated by Snail, Twist in the mesoderm specification network. Thirteen DRC genes were present in ≥7 TF_ALL datasets (DRC-13, Table S5), and had established functions in the development of mesodermal derivatives such as muscle, the nervous system and heart (Baylies and Bate, 1996; Bernard et al., 2010; Bray, 2016; Chen et al., 1996; Lo et al., 2002; Trujillo et al., 2016; Xie et al., 2016). Supporting NetNC-FTI predictions, *in situ* hybridisation for DRC-13 indicated earliest expression in (presumptive) mesoderm at: stages 4-6 (*wg, en, twi, N, htl, how*), stages 7-8 (*rib, pyd, mbc, abd-A*) and stages 9-10 (*pnt*) (BDGP; Hammonds et al., 2013; Hartley et al., 1987; Tomancak et al., 2002). The remaining two DRC-13 genes had no evidence for mesodermal expression (*fkh*) or no data available (*jar*). However, *fkh* is essential for caudal visceral mesoderm development (Kusch and Reuter, 1999) and *jar* is expressed in midgut mesoderm (Millo and Bownes, 2007).

Chromatin organisation clusters included several polycomb-group (PcG) and trithorax-group (TrxG) genes. Polycomb Repressive Complex 1 (PRC1) genes *ph-d*, *psc* (Shao et al., 1999) and *su(var)3-9* (Schotta et al., 2002) were identified most frequently in TF_ALL (Table S6). Further PcG/TrxG-related functional targets were: *ph-p* (Shao et al., 1999); c*orto* (Lopez et al., 2001); the TrxG-related gene *lolal* (Mishra et al., 2003); *taranis* (Schuster and Smith-Bolton, 2015); and TrxG genes *trithorax, moira* (Crosby et al., 1999; Tie et al., 2014). The gene silencing factor s*u(var)205* was also found in four TF_ALL datasets (Fanti et al., 1998). Therefore, NetNC predicted Snail and Twist regulation of: PRC1 core components and other gene silencing factors; TrxG genes; modifiers of PcG, TrxG activity. PcG genes are crucial oncofetal regulators and the focus of significant cancer drug development efforts (Koppens and Lohuizen, 2016; Sparmann and Lohuizen, 2006). Results align with previous reports that gene silencing in EMT involves PcG (Herranz et al., 2008; Koppens and Lohuizen, 2016), *Snai1* recruitment of Polycomb Repressive Complex 2 members (Herranz et al., 2008), and support a model where EMT TFs control the expression of their own coregulators.

Six TF_ALL datasets had brain development clusters. Snail regulation of neural genes is consistent with its repression of ectodermal (neural) genes in the prospective mesoderm (Gilmour et al., 2017; Leptin, 1991; Wieschaus and Nüsslein-Volhard, 2016). Additionally, Snail is important for neurogenesis in fly and mammals (Ashraf and Ip, 2001; Zander et al., 2014). Therefore, binding to neural functional modules might reflect potentiation for rapid activation in combination with other transcription factors within neural developmental trajectories (Nevil et al., 2017; Sandmann et al., 2007). NetNC results predict novel Twist functions, for example in activation or repression of mushroom body neuroblast proliferation factors *retinal homeobox, slender lobes*, and *taranis*. The mushroom body is a prominent structure in the fly brain, important for olfactory learning and memory (Caron et al., 2013). Twist is typically a transcriptional activator (Gilmour et al., 2017) although may contribute to Snail’s repressive activity (Lin et al., 2015). Indeed, Twist-related protein 1 repressed Cadherin-1 in breast cancers (Vesuna et al., 2008)

### Breast cancer subtype is characterised by differential expression of orthologous Snail and Twist functional targets

Integration of NetNC predictions, *Notch* screens and breast cancer transcriptomes enabled analysis of conserved molecular networks that orchestrate epithelial remodelling in development and cancers. NetNC-FTI Snail and Twist targets included known cancer genes and also predicted novel drivers (Figure 4, Figure S3, Tables S3, S5, S6). The fly genome is relatively tractable for network studies, while data availability (e.g. ChIP-chip, ChIP-seq, genetic screens) is enhanced by both considerable community resources and the relative ease of experimental manipulation (Mohr et al., 2014). Breast cancer intrinsic molecular subtypes with distinct clinical trajectories have been extensively validated and complement clinico-pathological parameters (Cejalvo et al., 2017; Sørlie et al., 2003). These subtypes are known as luminal-A, luminal-B, HER2-overexpressing, normal-like and basal-like. While more recent studies have classified more subtypes, for example identifying ten groups (Curtis et al., 2012), the five subtypes employed in our analysis had been widely used, extensively validated, exhibited clear differences in prognosis, overlapped with subgroups defined using standard clinical markers (*ESR1, HER2*), and aligned with distinct treatment pathways (Cejalvo et al., 2017; Sørlie et al., 2003). NetNC-FTI networks for all nine TF_ALL datasets overlapped with known cancer pathways, including significant enrichment for *Notch* modifiers (*q*<0.05). Aberrant activation of *Notch* orthologues in breast cancers had been demonstrated, and linked with EMT-like signalling, particularly for basal-like and claudin-low subtypes (Barnawi et al., 2016; Ingthorsson et al., 2016; Stylianou et al., 2006). We hypothesised that orthologous genes from Snail and Twist functional targets would stratify breast cancers into clinically meaningful groups.

### Unsupervised clustering with predicted functional targets recovers breast cancer intrinsic subtypes

Fifty-seven human orthologues (ORTHO-57) were NetNC-FTI functional targets in ≥4 TF_ALL datasets and represented within integrated gene expression microarray data for 2999 breast tumours (BrC_2999) (Moleirinho et al., 2013). Unsupervised clustering using ORTHO-57 stratified BrC_2999 by intrinsic molecular subtype (Figure 5). Clustering with NetNC results for individual Twist and Snail datasets also recovered the intrinsic breast cancer subtypes (Figure S4). Heatmap features were annotated according to the dendrogram structure and gene expression intensity (Figure 5). The datasets sna_2-4h_Toll^10b^, twi_2-4h_Toll^10b^ represent embryos formed entirely from mesodermal lineages (Zeitlinger et al., 2007) and, together, had significantly greater proportion of basal-like breast cancer genes than the combined sna_2-3h_union, twi_2-3h_union datasets (*p*<8.0×10^−4^); consistent with EMT characteristics of basal-like breast cancers (Sarrió et al., 2008). Basal-like tumours were characterised by *EN1* and *NOTCH1*, aligning with previous findings (Barnawi et al., 2016; Beltran et al., 2014; Stylianou et al., 2006). Notch signalling modulation is a promising area for cancer therapy (Ntziachristos et al., 2014) and orthologues of *Notch* modifiers identified in our analysis provide a pool of candidates that could potentially inform development of companion diagnostics or combination therapies for agents targeting the notch pathway in basal-like breast cancers. Elevated *ETV6* expression was also a feature of basal-like cancers, where copy number amplifications and recurrent gene fusions were previously reported (Adélaïde et al., 2007; Letessier et al., 2005). The Luminal A subtype (feature_LumA) had similarities with luminal B (feature_LumB_2_, ERBB3, MYO6) and normal-like (DOCK1, ERBB3, MYO6) tumours. High *BMPR1B* expression was a defining feature of the luminal A subtype, in agreement with oncogenic BMP signalling in luminal epithelia (Chapellier et al., 2015). *BMP2* may be pleiotropic, promoting EMT characteristics in some contexts (Ma et al., 2005; Ren and Dijke, 2017). *BMP2* expression was typically high in basal-like tumours and low in luminal cancers, aligning with *BMP2* upregulation as a feature of the EMT programme in basal-like cancers. Several genes were highly expressed in both feature_LumB_1_ and *ESR1* negative subtypes (feature_ERneg), including *ECT2, SNRPD1, SRSF2, CBX3*; our data suggest that these genes might contribute to the worse survival outcomes for luminal B relative to luminal A cancers. (Sørlie et al., 2001, 2003). *SNRPD1* and *SRSF2* function in splicing (Bermingham et al., 1995) which was linked to survival of multiple basal-like breast cancer cell lines (Chan et al., 2017). *CBX3* and *ECT2* were correlated with poor prognosis (Liang et al., 2017; Wang et al., 2018). These genes typically had low expression in Luminal A and normal-like subtypes. Feature_LoExp represents genes with low detection rates and tumours populating feature_LoExp are a mixture of subtypes, largely from a single study (Popovici et al., 2010). Key EMT genes (*SNAI2, TWIST1, QKI*) were assigned to the NL centroid and had highest relative expression in normal-like tumours (feature_NL, Figure 5). Feature_NL also included homeobox transcription factors (*HOXA9, MEIS2*) and a secreted cell migration guidance gene (*SLIT2*). Some genes had high expression in both normal-like and basal-like cancers, including *QKI*, which regulates circRNA formation in EMT (Conn et al., 2015), and the *FZD1* wnt/β-catenin receptor. Moreover, genes in feature_Bas and feature_NL clustered together, reflecting expression similarities between normal-like and basal-like subtypes. EMT may confer stem-like cell properties (DiMeo et al., 2009; Mani et al., 2008; Schmidt et al., 2015) and our results were consistent with dedifferentiation or arrested differentiation due to activation of an EMT-like programme in NL cancers. Previous work found stem cell markers in normal-like cancers (Sieuwerts et al., 2009; Sørlie et al., 2001) and *SNAI2* was critical for maintenance of mammary stem cells and linked with a stem-like signature in breast cancers (Guo et al., 2012; Lawson et al., 2015). Cell-compositional effects, associated with high stromal content in NL tumours (Prat and Perou, 2011), could also explain EMT-like molecular characteristics. We also found many differences between normal-like and basal-like cancer subtypes (Figure 4, Figure S3). Our results demonstrated that NetNC functional targets from fly mesoderm development capture clinically relevant molecular features of breast cancers and proposed novel candidate drivers of tumour progression.

**Figure 5.**
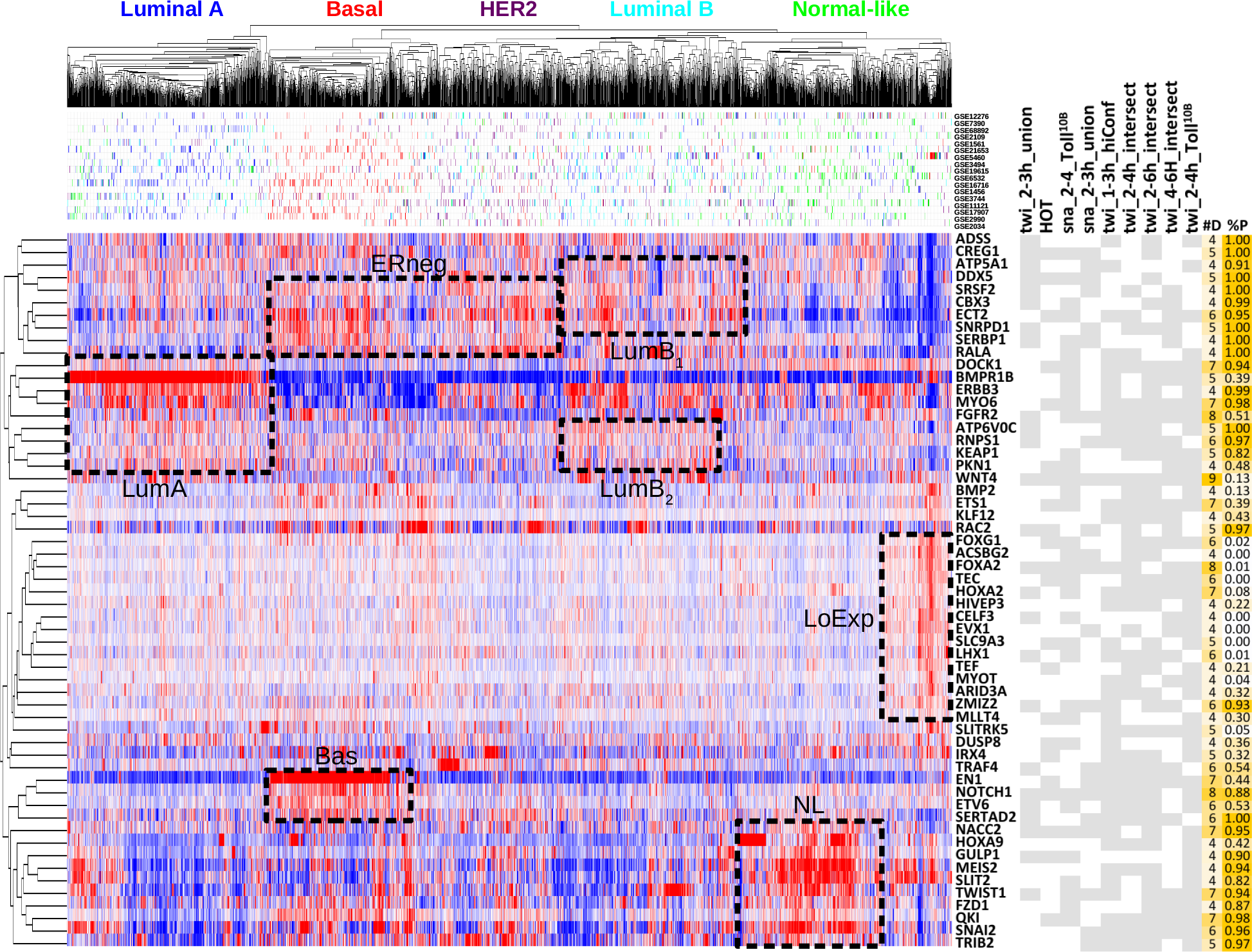
Predicted functional transcription factor targets capture human breast cancer biology. The heatmap shows results of unsupervised clustering with gene expression data for 2999 primary breast tumours and 57 orthologues of NetNC-FTI functional targets that were identified in at least four of nine TF_ALL datasets (ORTHO-57). Expression values were log2 transformed and mean-centred to give relative values across tumours (red=high, white=mean, blue=low). Intrinsic molecular subtype for each tumour is shown by the mosaic above the heatmap and below the dendogram, from left to right : luminal A (blue), basal-like (red), HER2-overexpressing (purple), luminal B (light blue) and normal-like (green). Source data identifiers are given to the right of the subtype mosaic. Features annotated onto the heatmap as black dashed lines identified genes upregulated in one or more intrinsic subtype; these features were termed ‘Bas’ (basal-like), ‘NL’(normal-like), ‘ERneg’ (basal-like and HER2-overexpressing), ‘LumB_1_’(luminal B), ‘LumB_2_’(luminal B), ‘LumA’ (luminal A) and ‘LoExp’ (low expression). The table to the right of the heatmap indicates inclusion (grey) or absence (white) of genes in NetNC-FTI results across the TF_ALL datasets. The column ‘#D’ gives the number of TF_ALL datasets where the gene was returned by NetNC-FTI and ‘%P’ column details the percentage of present calls for gene expression across the 2999 tumours. The LoExp feature corresponded overwhelmingly to genes with low %P values and to samples from a single dataset (Popovici et al., 2010). Some genes were annotated to more than one feature and reciprocal patterns of gene expression were found. For example, *BMPR1B, ERBB3* and *MYO6* were strongly upregulated in feature LumA but downregulated in basal-like and *HER2*-overexpressing cancers. Unexpectedly, feature NL (normal-like) had high expression of canonical EMT drivers, including *SNAI2*, *TWIST* and *QKI*. Some of the EMT genes in feature NL were also highly expressed in many basal-like tumours, while genes in feature Bas (*NOTCH, SERTAD2*) were upregulated in normal-like tumours. Also see Figure S4.

### Integrating NetNC functional target networks and breast cancer transcriptome profiling

Orthologous basal-like and normal-like genes were annotated onto NetNC-FTI networks, offering a new perspective on the molecular circuits controlling these different subtypes (Figure 4, Figure S3). We focussed on basal-like and normal-like cancers, which accounted for the large majority of networks studied and were prominent in centroid and heatmap analysis (Figure 5, Figure S4). Furthermore, EMT had been shown to be important for basal-like breast cancer biology (Sarrió et al., 2008) and, interestingly, key EMT genes were assigned to the normal-like subtype. Splicing factors and components of the ribosome were associated with the normal-like subtype in three networks (twi_2-4h_intersect, twi-2-6h_intersect, twi_2-3h_union); additionally, the proteasome and proteasome regulatory subunits in the twi_2-3h_union network had many genes annotated to the normal-like subtype.

The sna_2-4h_Toll^10b^ ‘RNA degradation and transcriptional regulation’ cluster was exclusively annotated to the basal-like subtype and included *HECA*, which may function as a tumour suppressor (Lin et al., 2013) and an oncogene (Chien et al., 2006). *HECA* was also identified in NetNC-FTI analysis of twi_2-4h_intersect and twi_4-6h_intersect; these datasets had Twist binding at different, non-contiguous sites that were both assigned to *hdc*, the fly HECA orthologue*. Hdc* was a multifunctional *Notch* signalling modifier, including in cell survival (Resende et al., 2017) and tracheal branching morphogenesis, activated by *escargot* (Steneberg et al., 1998). *HECA* was upregulated in basal-like relative to normal-like tumours (*p*<3.3×10^−23^). Taken together, these data support participation of *HECA* in an EMT-like gene expression programme in basal-like breast cancers.

The *SLC9A6* Na^+^/H^+^ antiporter was found in NetNC-FTI ion transport clusters for sna_2-4h_Toll^10b^ and twi_2-4h_Toll^10b^. Alterations in pH by Na^+^/H^+^ exchangers, particularly *SLC9A1*, drive basal-like breast cancer progression and chemoresistance (Amith and Fliegel, 2017). *SLC9A6* was 1.6-fold upregulated in basal-like relative to normal-like tumours (*p*<8.4×10^−71^) and might drive pH dysregulation as part of an EMT-like programme in basal-like cancers. Sna_2-4h_Toll^10b^ clusters were depleted in normal-like annotations, compared with Twist; 4/18 clusters had two or more normal-like orthologues, significantly fewer than twi_ 2-4h_Toll^10b^ (8/17, *p<*0.035) and twi_2-3h_union (12/27, binomial *p<*0.01). A further basal-like cluster for twi_2-3h_union was annotated to ‘mitochondrial translation’ (MT), an emerging area of interest for cancer therapy (Weinberg and Chandel, 2015). Our results highlight MT as a potentially attractive target in basal-like breast cancers, aligning with work linking MT upregulation with loss of *RB1* and *p53* (Jones et al., 2016).

Chromatin organisation clusters frequently associated with basal-like annotations. For example, the twi_2-3h_union ‘chromatin organisation and transcriptional regulation’ cluster had six basal-like genes, including three Notch modifiers (*ash1*, *tara, Bap111).* These were orthologous to the *ASH1L* histone methyltransferase and candidate poor prognosis factor with copy number amplifications in basal-like tumours (Liu et al., 2014); the *SERTAD2* bromodomain interacting oncogene and E2F activator (Cheong et al., 2009); and *SMARCE1*, a core subunit of the SWI/SNF chromatin remodelling complex that regulated *ESR1*, interacted with *HIF1A* signalling and potentiated breast cancer metastasis (García-Pedrero et al., 2006; Sethuraman et al., 2016; Sokol et al., 2017). Notch can promote EMT-like characteristics and mediated hypoxia-induced invasion in multiple cell lines (Sahlgren et al., 2008). Consistent with these studies, our work supported conserved function for *SMARCE1* in EMT-like signalling, both in mesoderm development and basal-like breast cancers, possibly downstream of *NOTCH1* and through regulation of SWI/SNF targeting. Indeed, SWI/SNF controled chromatin switching in oral cancer EMT (Mohd-Sarip et al., 2017). *Taranis*, orthologous to *SERTAD2*, also functions to stabilise the expression of *engrailed* in regenerating tissue (Schuster and Smith-Bolton, 2015). The *engrailed* orthologue *EN1* was the clearest single basal-like biomarker in the data examined (Figure 5) and acts as a survival factor (Beltran et al., 2014). *SERTAD2* could cooperate with *EN1* in basal-like cancers; our results evidence coordinated expression of these two genes as part of a gene expression programme controlled by EMT TFs. Regulation of *EN1, SERTAD2* within an EMT programme could harmonise previous results demonstrating key roles for both neural-specific and EMT TFs in basal-like breast cancers (Beltran et al., 2014; Sarrió et al., 2008). Therefore, we identify chromatin organisation factors downstream of Snail and Twist with orthologues that may control *Notch* output and breast cancer progression through a chromatin remodelling mechanism. Indeed, NetNC results predict components of feedback loops where EMT TFs regulate chromatin organisation genes that, in turn, may both reinforce and coordinate downstream stages of gene expression programmes for mesoderm development and cancer progression. Stages of the EMT programme were described elsewhere, reviewed in (Nieto et al., 2016); our results map networks that may control the remodelling of Waddington’s landscape - identifying crosstalk between Snail, Twist, epigenetic modifiers and regulation of key developmental pathways (Hemberger et al., 2009). Dynamic interplay between successive cohorts of TFs and chromatin organisation factors could be an attractive mechanism to determine progress through and the ordering of steps in (partial) EMTs, consistent with ‘metastable’ intermediate stages (Nieto et al., 2016).

### Novel Twist and Snail functional targets influence invasion in a breast cancer model of EMT

NetNC results predicted new gene functions in EMT and cell invasion, including *SNX29* (also known as *RUNDC2A*), *ATG3*, *IRX4* and *UNK*. We investigated the functional and instructive role of these genes in an established invasion model (Dhasarathy et al., 2007). MCF7 cells are weakly invasive (Lacroix and Leclercq, 2004), thus the *SNAI1-*inducible MCF7 cell line was well suited to study alteration in expression of the selected genes in terms of their influence on invasion in conjunction with *SNAI1* induction, knockdown or independently (Figure 6). Potential artefacts associated with changes in cell growth or proliferation are controlled within the transwell assays used, because values reflect the ratio of signal from cells located at either side of the matrigel barrier.

**Figure 6.**
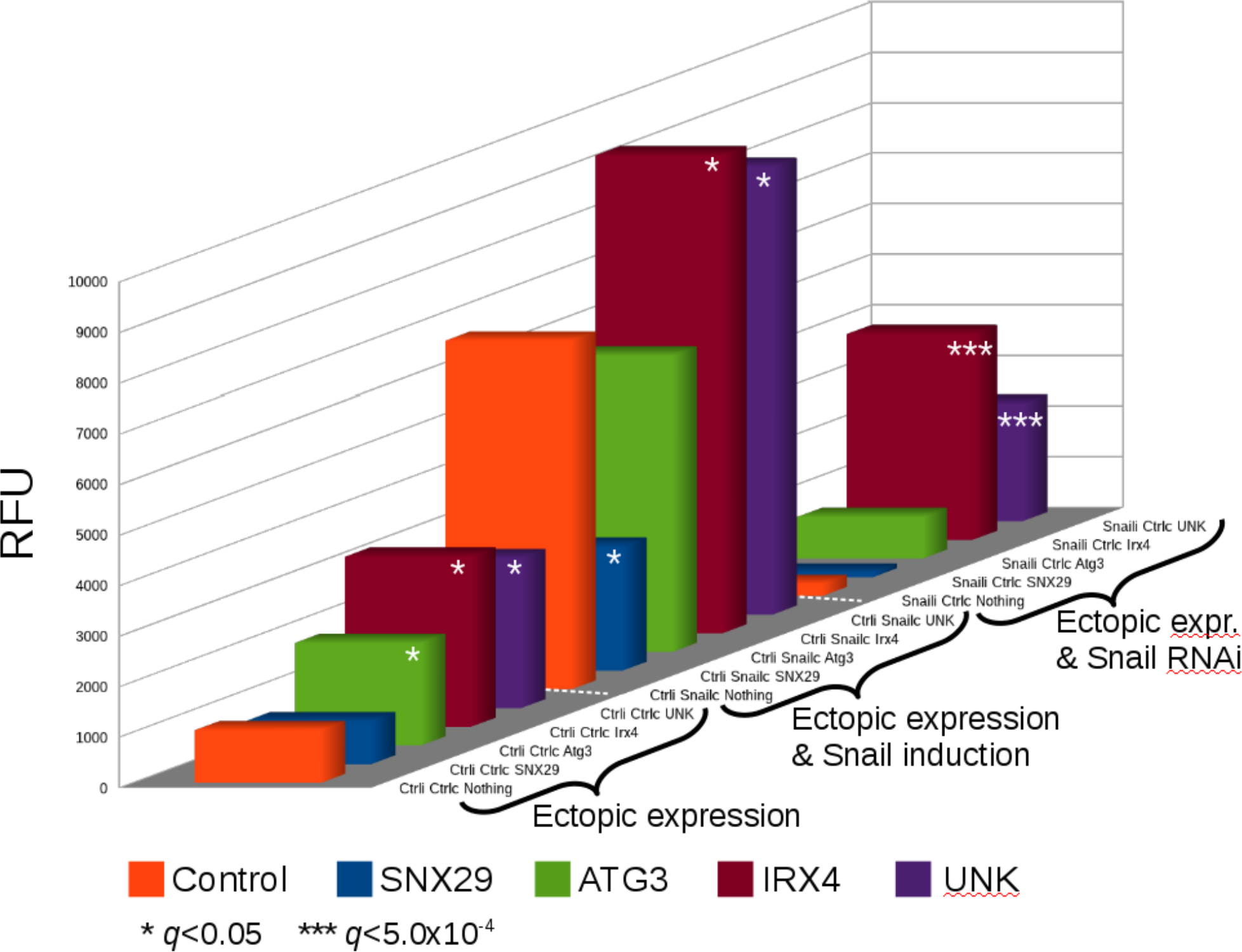
Validation of candidate invasion genes in breast cancer cells. Fluorescence from invasive MCF7 cells is shown. Induction of every gene examined significantly changed invasion in least one of three conditions: a) ectopic expression; b) ectopic expression and *SNAI1* induction; c) ectopic expression with shRNA knockdown of *SNAI1*. *SNX29* (blue) had reduced invasion compared with the *SNAI1* induction control (orange); *UNK* (purple) and *IRX4* (dark red) had increased invasion in all three conditions examined; *ATG3* had higher invasion at background levels of *SNAI1* (without induction or knockdown). Mean values are shown, n=3 datapoint; statistical significance is shown by * *q*<0.05; *** *q*<5.0×10^−4^. Also see Supplemental File 2.

Over-expression of *IRX4* significantly increased invasion relative to controls in all conditions examined and *IRX4* had high relative expression in a subset of basal-like breast cancers (Figures 5, 6). *IRX4* is a homeobox transcription factor involved in cardiogenesis, marking a ventricular-specific progenitor cell (Nelson et al., 2016) and is also associated with prostate cancer risk (Xu et al., 2014). *SNX29* belongs to the sorting nexin protein family that function in endosomal sorting and signalling (Marat and Haucke, 2016). *SNX29* is poorly characterised and ectopic expression significantly reduced invasion in a *SNAI1*-dependent manner (Figure 6). Since we obtained these results, *SNX29* downregulation was associated with metastasis and chemoresistance in ovarian carcinoma (Zhu et al., 2015), consistent with *SNX29* inhibition of invasion driven by Snail. *ATG3* is an E2-like enzyme required for autophagy and mitochondrial homeostasis (Doherty and Baehrecke, 2018), *ATG3* overexpression significantly increased MCF7 invasion. Knockdown of *ATG3* reduced invasion in hepatocellular carcinoma (Li et al., 2013). *UNK* is a RING finger protein homologous to *unkempt* which binds mRNA, functions in ubiquitination and was upregulated in gastrulation (Mohler et al., 1992). Others reported that *UNK* mRNA binding controls neuronal morphology and can induce spindle-like cell shape in fibroblasts (Murn et al., 2015, 2016). *UNK* significantly increased MCF7 invasion both independently of and additively with Snail; supporting a potential role in breast cancer progression. Indeed, *UNK* was overexpressed in cancers relative to controls in ArrayExpress (Parkinson et al., 2009). These *in vitro* confirmatory results both support the novel analysis approach and evidence new function for the genes examined.

## METHODS

### A Comprehensive *D. melanogaster* functional gene network (DroFN)

A high-confidence, comprehensive *Drosophila melanogaster* functional network (DroFN) was developed using a previously described inference approach (Overton et al., 2011). Functional interaction probabilities, corresponding to pathway co-membership, were estimated by logistic regression of Bayesian probabilities from STRING v8.0 scores (Jensen et al., 2009) and Gene Ontology (GO) coannotations (Ashburner et al., 2000), taking KEGG (Kanehisa et al., 2010) pathways as gold standard.

Gene pair co-annotations were derived from the GO database of March 25th 2010. The GO Biological Process (BP) and Cellular Component (CC) branches were read as a directed graph and genes added as leaf terms. The deepest term in the GO tree was selected for each gene pair, and BP was given precedence over CC. Training data were taken from KEGG v47, comprising 110 pathways (TRAIN-NET). Bayesian probabilities for STRING and GO coannotation frequencies were derived from TRAIN-NET (Overton et al., 2011). Selection of negative pairs from TRAIN-NET using the *perl* rand() function was used to generate training data with equal numbers of positive and negative pairs (TRAIN-BAL), which was input for logistic regression, to derive a model of gene pair functional interaction probability:

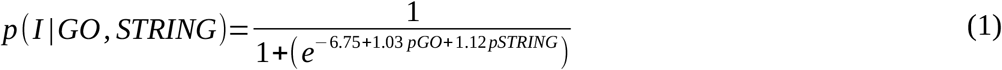

Where:

pGO is the Bayesian probability derived from Gene Ontology coannotation frequency pSTRING is the Bayesian probability derived from the STRING score frequency

The above model was applied to TRAIN-NET and the resulting score distribution thresholded by seeking a value that maximised the F-measure (van Rijsbergen, 1979) and True Positive Rate (TPR), while also minimising the False Positive Rate (FPR). The selected threshold value (*p ≥*0.779) was applied to functional interaction probabilities for all possible gene pairs to generate the high-confidence network, DroFN.

For evaluation of the DroFN network, time separated test data (TEST-TS) were taken from KEGG v62 on 13/6/12, consisting of 14 pathways that were not in TRAIN-NET. TEST-TS was screened against TRAIN-NET, eliminating 34 positive and 218 negative gene pairs to generate the blind test dataset TEST-NET (4599 pairs). GeneMania (version of 10^th^ August 2011) (Warde-Farley et al., 2010) and DROID (v2011_08) (Yu et al., 2008) were assessed against TEST-NET (Table S1, Figure S1).

Enrichment of DroFN edges in DroPIM was estimated as follows: a total of 999 genes were found in both DroFN and the DroPIM network thresholded at FDR 0.05 (DroPIM_FDR). These 999 genes had 5747 edges in DroPIM_FDR and 25797 edges in DroFN, of which 2175 were common to both networks. A 2×2 contingency table was constructed conditioning on the presence of edges for these 999 genes in the DroFN and DroPIM_FDR networks. The contingency table cell corresponding to edges not found in DroFN or DroPIM was populated by the number of possible edges for the 999 genes ((n^2^ - n) / 2), subtracting the values from the other cells. Therefore the contingency table cell values were: 2175, 3572, 23622, 469132. The enrichment *p*-value was calculated by Fisher’s Exact Test.

### Network neighbourhood clustering (NetNC) algorithm

NetNC identifies functionally coherent nodes in a subgraph *S* of functional gene network *G* (an undirected graph), induced by some set of nodes of interest *D*; for example, candidate transcription factor target genes assigned from analysis of ChIP-seq data. Intuitively, we consider the proportion of common neighbours for nodes in *S* to define coherence; for example, nodes that share neighbours have greater coherence than nodes that do not share neighbours. The NetNC workflow is summarised in Figure 1 and described in detail below. Two analysis modes are available a) node-centric (parameter-free) and b) edge-centric, with two parameters. Both modes begin by assigning a p-value to each edge (*S*_*ij*_) from Hypergeometric Mutual Clustering (HMC) (Goldberg and Roth, 2003), described in points one and two, below.

1. A two times two contingency table is derived for each edge *S*_*ij*_ by conditioning on the Boolean connectivity of nodes in *S* to *S*_*i*_ and *S*_*j*_. Nodes *S*_*i*_ and *S*_*j*_ are not counted in the contingency table.
2. Exact hypergeometric *p*-values (Goldberg and Roth, 2003) for enrichment of the nodes in *S* that have edges to the nodes *S*_*i*_ and *S*_*j*_ are calculated using Fisher’s Exact Test from the contingency table. Therefore, a distribution of p-values (*H*_1_) is generated for all edges *S*_*ij*_.
3. The NetNC edge-centric mode employs positive false discovery rate (Storey, 2002) and an iterative minimum cut procedure (Ford and Fulkerson, 1956) to derive clusters as follows:
  a. Subgraphs with the same number of nodes as *S* are resampled from *G*, application of steps 1 and 2 to these subgraphs generates an empirical null distribution of neighbourhood clustering *p*-values (*H*_0_). This *H*_0_ accounts for the effect of the sample size and the structure of *G* on the *S*_*ij*_ hypergeometric *p*-values (*p*_*ij*_). Each NetNC run on TF_ALL in this study resampled 1000 subgraphs to derive *H*_0_.
  b. Each edge in *S* is associated with a positive false discovery rate (*q*) estimated over *p*_*ij*_ using *H*_1_ and *H*_0_. The neighbourhood clustering subgraph *C* is induced by edges where the associated *q* ≤ *Q*.
  c. An iterative minimum cut procedure (Ford and Fulkerson, 1956) is applied to *C* until all components have density greater than or equal to a threshold *Z*. Edge weights in this procedure are taken as the negative log *p*-values from *H*_1_.
  d. As described in section 4.2.3, thresholds *Q* and *Z* were chosen to optimise the performance of NetNC on the ‘Functional Target Identification’ task using training data taken from KEGG. Connected components with less than three nodes are discarded, in line with common definitions of a ‘cluster’. Remaining nodes are classified as functionally coherent.
4. The node-centric, parameter-free mode proceeds by calculating degree-normalised node functional coherence scores (NFCS) from *H*_1_, then identifies modes of the NFCS distribution using Gaussian Mixture Modelling (GMM) (Lubbock et al., 2013):
  a. The node functional coherence score (NFCS) is calculated by summation of *S*_*ij*_ *p*-values in *H*_1_ (*p*_*ij*_) for fixed *S*_*i*_, normalised by the *S*_*i*_ degree value in *S* (*di*):

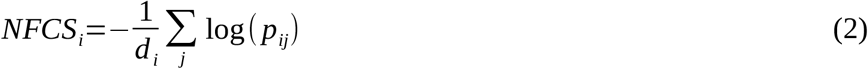
  b. GMM is applied to identify structure in the NFCS distribution. Expectation-maximization fits a mixture of Gaussians to the distribution using independent mean and standard deviation parameters for each Gaussian (Dempster et al., 1977; Lubbock et al., 2013). Models with 1..9 Gaussians are fitted and the final model selected using the Bayesian Information Criterion (BIC).
  c. Nodes in high-scoring mode(s) are predicted to be ‘Functionally Bound Targets’ (FBTs) and retained. Firstly, any mode at NFCS<0.05 is excluded because this typically represents nodes with no edges in *S* (where NFCS=0). A second step eliminates the lowest scoring mode if >1 mode remains. Very rarely a unimodal model is returned, which may be due to a large non-Gaussian peak at NFCS=0 confounding model fitting; if necessary this is addressed by introducing a tiny Gaussian noise component (SD=0.01) to the NFCS=0 nodes to produce NFCS_GN0. GMM is performed on NFCS_GN0 and nodes eliminated according to the above procedure on the resulting model. This procedure was developed following manual inspection of results on training data from KEGG pathways with ‘synthetic neutral target genes’ (STNGs) as nodes resampled from *G* (TRAIN-CL, described in section 2.2.1).

Therefore, NetNC can be applied to predict functional coherence using either edge-centric or node-centric analysis modes. The edge-centric mode automatically produces a network, whereas the node-centric analysis does not output edges; therefore to generate networks from predicted FBT nodes an edge pFDR threshold may be applied, pFDR≤0.1 was selected as the default value. The statistical approach to estimate pFDR and local FDR are described in the sections below.

### Estimating positive false discovery rate for hypergeometric mutual clustering p-values

The following procedure is employed to estimate positive False Discovery Rate (pFDR) (Storey, 2002) in the NetNC edge-centric mode. Subgraphs with number of nodes identical to *S* are resampled from *G* to derive a null distribution of HMC *p*-values (*H_0_*) (section 4.2, above). The resampling approach for pFDR calculation in NetNC-FTI controls for the structure of the network *G*, including degree distribution, but does not control for the degree distribution or other network properties of the subgraph *S* induced by the input nodelist (*D*). In scale free and hierarchical networks, degree correlates with clustering coefficient; indeed, this property is typical of biological networks (Yamada and Bork, 2009). Part of the rationale for NetNC assumes that differences between the properties of *G* and *S* (for example; degree, clustering coefficient distributions) may enable identification of clusters within *S*. Therefore, it would be undesirable to control for the degree distribution of *S* during the resampling procedure for pFDR calculation because this would also partially control for clustering coefficient. Indeed clustering coefficient is a node-centric parameter that has similarity with the edge-centric Hypergeometric Clustering Coefficient (HMC) calculation (Goldberg and Roth, 2003) used in the NetNC algorithm to analyse *S*. Hence, the resampling procedure does not model the degree distribution of *S*, although the degree distribution of *G* is controlled for. Positive false discovery rate is estimated over the *p*-values in *H*_1_ (*p*_*ij*_) according to Storey (Storey, 2002):

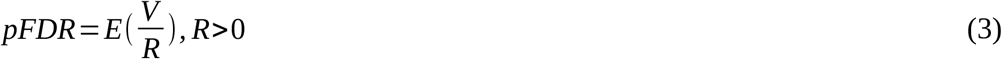

Where:

*R* denotes hypotheses (edges) taken as significant

*V* are the number of false positive results (type I error)

NetNC steps through threshold values (*p*_*α*_) in *p*_*ij*_ estimating *V* using edges in *H*_0_ with *p*≤*p*_*α*_. *H*_0_ represents *Y* resamples, therefore *V* is calculated at each step:

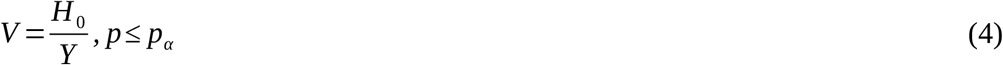

The *H*_1_ *p*-value distribution is assumed to include both true positives and false positives (FP); *H*_0_ is taken to be representative of the FP present in *H*_1_. This approach has been successfully applied to peptide spectrum matching (Fitzgibbon et al., 2008; Sennels et al., 2009). The value of *R* is estimated by:

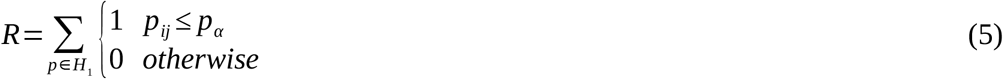

Additionally, there is a requirement for monotonicity:

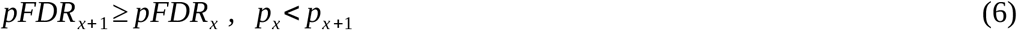

Equation (6) represents a conservative procedure to prevent inconsistent scaling of pFDR due to sampling effects. For example consider the scaling of pFDR for pFDR_x+1_ at a *p*_*ij*_ value with additional edges from *H*_1_ but where no more resampled edges (i.e. from *H*_0_) were observed in the interval between p_x_ and p_x+1_; before application of equation (6), the value of pFDR_x+1_ would be lower than pFDR_x_. The approach also requires setting a maximum on estimated pFDR, considering that there may be values of *p*_*α*_ where *R* is less than *V*. We set the maximum to 1, which would correspond to a prediction that all edges at *p*_*ij*_ are FPs. The assumption that *H*_1_ includes false positives is expected to hold in the context of candidate transcription factor target genes and also generally across biomedical data due to the stochastic nature of biological systems (Marusyk et al., 2012; Raj and van Oudenaarden, 2008; Raj et al., 2010). We note that an alternative method to calculate R using both *H*_1_ and *H*_0_ would be less conservative than the approach presented here.

### Estimating local false discovery rate from global false discovery rate

We developed an approach to estimate local false discovery rate (lcFDR) (Efron et al., 2001), being the probability that an object at a threshold (*p*_α_) is a false positive (FP). Our approach takes global pFDR values as basis for lcFDR estimation. In the context of NetNC analysis using the DroFN network, a FP is defined as a gene (node) without a pathway comembership relationship to any other nodes in the nodelist *D*. The most significant pFDR value (pFDR_min_) from NetNC was determined for each node *S*_*i*_ across the edge set *S*_*ij*_. Therefore, pFDR_min_ is the pFDR value at which node *S*_*i*_ would be included in a thresholded network. We formulated lcFDR for the nodes with pFDR_min_ meeting a given *p*_*α*_ (*k*) as follows:

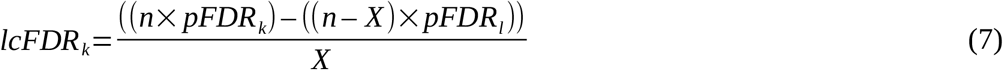

Where *l* denotes the pFDR_min_ closest to and smaller than *k*, and where at least one node has pFDR_min_≡pFDR_l_. Therefore, our approach can be conceptualised as operating on ordered pFDR_min_ values. *n* indicates the nodes in *D* with pFDR_min_ values meeting threshold *k. X* represents the number of nodes at *p*_*α*_ ≡*k.* The number of FPs for nodes with *p*_*α*_ ≡*k* (FP_*k*_) is estimated by subtracting the FP for threshold *l* from the FP at threshold *k.* Thus, division of FP_*k*_ by *X* gives local false discovery rate bounded by *k* and *l* (Figure S5). If we define the difference between pFDR_*k*_ and pFDR_*l*_:

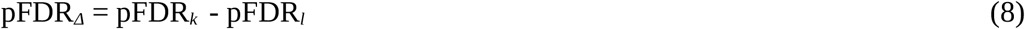

Substituting pFDR_*k*_ for (pFDR_*l*_ + pFDR_*Δ*_) into equation (7) and then simplifying gives:

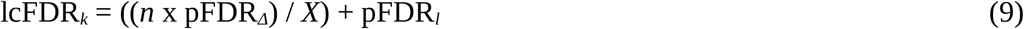

Equations (7) and (9) do not apply to the node(s) in *D* at the smallest possible value of pFDR_min_ because pFDR_l_ would be undefined; instead, the value of lcFDR_*k*_ is calculated as the (global) pFDR_min_ value. Indeed, global FDR and local FDR are equivalent when *H*_1_ consists of objects at a single pFDR_min_ value. Taking the mean lcFDR_*k*_ across *D* provided an estimate of neutral binding in the studied ChIP-chip, ChIP-seq datasets and was calibrated against mean lcFDR values from datasets that had a known proportion of Synthetic Neutral Target Genes (SNTGs). Estimation of the total proportion of neutral binding in ChIP-chip or ChIP-seq data required lcFDR rather than (global) pFDR and, for example, accounts for the shape of the *H*_1_ distribution. In the context of NetNC analysis of TF_ALL, mean lcFDR may be interpreted as the probability that any candidate target gene is neutrally bound in the dataset analysed; therefore providing estimation of the total neutral binding proportion. Computer code for calculation of lcFDR is provided within the NetNC distribution. Estimates of SNTGs by the NetNC-FBT approach were not taken forward due to large 95% CI values (Figure S8).

### Median difference and correlation between estimates of neutral binding from NetNC Functional Target Identification and local False Discovery Rate

Candidate target genes that did not pass NetNC-FTI thresholds were labelled neutrally bound (FTI_NB). The proportion of FTI_NB genes was compared to the proportion of neutral binding estimated by lcFDR (lcFDR_NB, Figure 3A). The modulus of the difference between FTI_NB and lcFDR_NB for each dataset gave a distribution of differences in neutral binding proportion and the median of this distribution was quoted in the text above.

### NetNC benchmarking and parameter optimisation

Gold standard data for NetNC benchmarking and parameterisation were taken as pathways from KEGG (v62, downloaded 13/6/12) (Kanehisa et al., 2010). Training data were selected as seven pathways (TRAIN-CL, 184 genes) and a further eight pathways were selected as a blind test dataset (TEST-CL, 186 genes) summarised in Table S7. For both TRAIN-CL and TEST-CL, pathways were selected to be disjoint and to cover a range of different biological functions. However, pathways with shared biology were present within each group; for example TRAIN-CL included the pathways dme04330 ‘Notch signaling’ and dme04914 ‘Progesterone-mediated oocyte maturation’, which are related by notch involvement in oogenesis (López-Schier and St Johnston, 2001; Schmitt and Nebreda, 2002). TEST-CL also included the related pathways dme04745 ‘Phototransduction’ and dme00600 ‘Sphingolipid metabolism’, for example where ceramide kinase regulates photoreceptor homeostasis (Acharya et al., 2003; Dasgupta et al., 2009; Yonamine et al., 2011).

Gold standard datasets were also developed in order to investigate the effect of dataset size and noise on NetNC performance. The inclusion of noise as resampled network nodes into the gold-standard data was taken to model neutral TF binding (Li et al., 2008; Shlyueva et al., 2014) and matches expectations on data taken from biological systems in general (Marusyk et al., 2012; Raj and van Oudenaarden, 2008). Therefore, gold standard datasets were generated by combining TRAIN-CL with nodes resampled from the network (*G*) and combining these with TRAIN-CL. The final proportion of resampled nodes (Synthetic Neutral Target Genes, SNTGs) ranged from 5% through to 80% in 5% increments. Since we expected variability in the network proximity of SNTGs to pathway nodes (*S*), 100 resampled datasets were generated per %SNTG increment. Further gold-standard datasets were generated by taking five subsets of TRAIN-CL, from three through seven pathways. Resampling was applied for these datasets as described above to generate node lists representing five pathway sets in TRAIN-CL by sixteen %SNTG levels by l00 repeats (TRAIN_CL_ALL, 8000 node lists). A similar procedure was applied to TEST-CL, taking from three through eight pathways to generate data representing six pathway subsets by sixteen noise levels by 100 repeats (TEST-CL_ALL, 9600 node lists). Data based on eight pathways (TEST-CL_8PW, 1600 node lists) were used for calibration of lcFDR estimates. Preliminary training and testing against the MCL algorithm (Enright et al., 2002) utilised a single subsample for 10%, 25%, 50% and 75% SNTGs (TRAIN-CL-SR, TEST-CL-SR).

NetNC analysed the TRAIN-CL_ALL datasets in edge-centric mode, across a range of FDR (*Q*) and density (*Z*) threshold values. Performance was benchmarked on the Functional Target Identification (FTI) task which assessed the recovery of biological pathways and exclusion of SNTGs. Matthews correlation coefficient (MCC) was computed as a function of NetNC parameters (Q, Z). MCC is attractive because it is captures predictive power in both the positive and negative classes. FTI was a binary classification task for discrimination of pathway nodes from noise, therefore all pathway nodes were taken as as positives and SNTGs were negatives for the FTI MCC calculation. The FTI approach therefore tests discrimination of pathway nodes from SNTGs, which is particularly relevant to identification of functionally coherent candidate TF targets from ChIP-chip or ChIP-seq peaks.

Parameter selection for NetNC on the FTI task analysed MCC values for the 100 SNTG resamples across five pathway subsets by sixteen SNTG levels in TRAIN-CL_ALL over the Q, Z values examined, respectively ranging from up to 10^−7^ to 0.8 and from up to 0.05 to 0.9. Data used for optimisation of NetNC parameters (Q, Z) are available from the Biomodels database and contour plots showing mean MCC across Q, Z values per %SNTG are provided in Figure S7. A ‘SNTG specified’ parameter set was developed for situations where an estimate of the input data noise component is available, for example from the node-centric mode of NetNC. In this parameterisation, for each of the sixteen datasets with different proportions of SNTG (5%.. 80%), MCC values were normalized across the five pathway subsets of TRAIN-CL (from three through seven pathways), by setting the maximum MCC value to 1 and scaling all other MCC values accordingly. The normalised MCC values <0.75 were set to zero and then a mean value was calculated for each %SNTG value across five pathway subsets by 100 resamples in TRAIN-CL_ALL (500 datasets per noise proportion). This approach therefore only included parameter values corresponding to MCC performance ≥75% of the maximum across the five TRAIN-CL pathway subsets. The high performing regions of these ‘summary’ contour plots sometimes had narrow projections or small fragments, which could lead to parameter estimates that do not generalise well on unseen data. Therefore, parameter values were selected as the point at the centre of the largest circle (in (Q, Z) space) completely contained in a region where the normalised MCC value was ≥0.95. This procedure yielded a parameter map: (SNTG Estimate) → (Q, Z), given in Table S8. NetNC parameters were also determined for analysis without any prior belief about the %SNTG in the input data - and therefore generalise across a wide range of %SNTG and dataset sizes. For this purpose, a contour plot was produced to represent the proportion of datasets where NetNC performed better than 75% of the maximum performance across TRAIN-CL_ALL for the FTI task in the Q, Z parameter space. The maximum circle approach described above was applied to the contour plot in order to derive ‘robust’ parameter values (Q, Z), which were respectively 0.120, 0.306 (NetNC-FTI).

### Performance on blind test data

We compared NetNC against leading methods, HC-PIN (Wang et al., 2011) and MCL (Enright et al., 2002) on blind test data (Figure 2, Table S2). Previous work that evaluated nine clustering algorithms, including MCL, found that HC-PIN had strong performance in functional module identification and was robust against false positives (Wang et al., 2011); therefore HC-PIN was selected for extensive comparison against NetNC. Input, output and performance summary files for HC-PIN on TEST-CL are available from the Biomodels database (per datapoint, n=100 for NetNC, n=99 for HC-PIN). HC-PIN was run on the weighted graphs induced in DroFN by TEST-CL with default parameters (lambda = 1.0, threshold size = 3). MCL clusters in DroFN significantly enriched for query nodes from TEST-CL-SR were identified by resampling to generate a null distribution (Overton et al., 2011). Clusters with *q*<0.05 were taken as significant. MCL performance was optimised for the Functional Target Identification (FTI) task over the TRAIN-CL-SR datasets for MCL inflation values from 2 to 5 incrementing by 0.2. The best-performing MCL inflation value overall was 3.6 (Table S9).

### Subsampling of transcription factor binding datasets and statistical testing

Robustness of NetNC performance was studied by taking 95%, 80% and 50% resamples from nine public transcription factor binding datasets, summarised in section 4.3 and described previously in detail (MacArthur et al., 2009; Ozdemir et al., 2011; Roy et al., 2010; Sandmann et al., 2007; Zeitlinger et al., 2007). A hundred subsamples of each of these datasets were taken at rates of 95%, 80% and 50%, thereby producing a total of 2700 datasets (TF_SAMPL). NetNC-FTI results across TF_SAMPL were used as input for calculation of median and 95% confidence intervals for the edge and gene overlap per subsampling rate for each transcription factor dataset analysed. The NetNC resampling parameter (Y) was set at 100, the default value. The edge overlap was calculated as the proportion of edges returned by NetNC-FTI for the subsampled dataset that were also present in NetNC-FTI results for the full dataset (i.e. at 100%). Therefore, nine values for median overlap and 95% CI were produced per subsampling rate for both edge and gene overlap, corresponding to the nine transcription factor binding datasets (Table S4). The average (median) value of these nine median overlap values, and of the 95% CI, was calculated per subsampling rate; these average values are quoted in Supplemental Material.

False discovery rate (FDR) correction of *p*-values was applied where appropriate and is indicated in this manuscript by the commonly used notation ‘*q’* Benjamini-Hochberg correction was applied (Benjamini and Hochberg, 1995) unless otherwise specified in the text. Calculation of pFDR and local FDR values by NetNC is described in the sections above.

### Transcription factor binding and Notch modifier datasets

We analysed public Chromatin Immunoprecipitation (ChIP) data for the transcription factors *twist* and *snail* in early *Drosophila melanogaster* embryos. These datasets were derived using ChIP followed by microarray (ChIP-chip) (MacArthur et al., 2009; Sandmann et al., 2007; Zeitlinger et al., 2007) and ChIP followed by solexa pyrosequencing (ChIP-seq) (Ozdemir et al., 2011). Additionally ‘highly occupied target’ regions, reflecting multiple and complex transcription factor occupancy profiles, were obtained from ModEncode (Roy et al., 2010). Nine datasets were analysed in total (TF_ALL) and are summarised below.

The ‘union’ datasets (WT embryos 2-3h, mostly late stage four or early stage five) combined ChIP-chip peaks significant at 1% FDR for two different antibodies targeted at the same TF and these were assigned to the closest transcribed gene according to RNA Polymerase II binding data (MacArthur et al., 2009). Additionally, where the closest transcribed gene was absent from the DroFN network then the nearest gene was included if it was contained in DroFN. This approach generated the datasets sna_2-3h_union (1158 genes) and twi_2-3h_union (1848 genes). The union of peaks derived from two separate antibodies maximised sensitivity and may have reduced potential false negatives arising from epitope steric occlusion. For the ‘Toll^10b^’ datasets, significant peaks with at least two-fold enrichment for Twist or Snail binding were taken from ChIP-chip data on Toll^10b^ mutant embryos (2-4h), which had constitutively activated Toll receptor (Stathopoulos et al., 2002; Zeitlinger et al., 2007); mapping to DroFN generated the datasets twi_2-4h_Toll^10b^ (1238 genes), sna_2-4h_Toll^10b^ (1488 genes). Toll^10b^ embryos had high expression of Snail and Twist, which drove all cells to mesodermal fate trajectories (Zeitlinger et al., 2007). The two-fold enrichment threshold selected for this study reflects ‘weak’ binding, although was expected to include functional TF targets (Biggin, 2011). Therefore the candidate target genes for twi_2-4h_Toll^10b^ and sna_2-4h_Toll^10b^ were expected to contain a significant proportion of false positives. The Highly Occupied Target dataset included 38562 regions, of which 1855 had complexity score ≥8 and had been mapped to 1648 FlyBase genes according to the nearest transcription start site (Roy et al., 2010); 677 of these genes were matched to a DroFN node (HOT). The ‘HighConf’ data took Twist ChIP-seq binding peaks in WT embryos (1-3h) that had been reported to be ‘high confidence’ assignments; high confidence filtering was based on overlap with ChIP-chip regions, identification by two peak-calling algorithms and calibration against peak intensities for known Twist targets, corresponding to 832 genes (Ozdemir et al., 2011). A total of 664 of these genes were found in DroFN (twi_1-3h_hiConf) and represented the most stringent approach to peak calling of all the nine TF_ALL datasets. The intersection of ChIP-chip binding for two different Twist antibodies in WT embryos spanning two time periods (2-4h and 4-6h) identified a total of 1842 target genes (Sandmann et al., 2007) of which 1444 mapped to DroFN (Intersect_ALL). Subsets of Intersect_ALL identified regions bound only at 2-4 hours (twi_2-4h_intersect, 801 genes), or only at 4-6 hours (twi_4-6h_intersect, 818 genes), or ‘continuously bound’ regions identified at both 2-4 and 4-6 hours (twi_2-6h_intersect, 615 genes). Assigned gene targets may belong to more than one subset of Intersect_ALL because time-restricted binding was assessed for putative enhancer regions prior to gene mapping; overlap of the Intersect_ALL subsets ranged between 30.2% and 55.4%. The Intersect_ALL datasets therefore enabled assessment of functional enhancer binding according to occupancy at differing time intervals and also to examine the effect of intersecting ChIPs for two different antibodies upon the proportion of predicted functional targets recovered.

Seven of the nine TF_ALL datasets included developmental time periods encompassing stage four (syncytial blastoderm, 80-130 minutes), cellularisation of the blastoderm (stage five, 130-170 minutes) and initiation of gastrulation (stage 6, 170-180 minutes) (Campos-Ortega and Hartenstein, 1997; MacArthur et al., 2009; Ozdemir et al., 2011; Sandmann et al., 2007; Zeitlinger et al., 2007). The datasets twi_2-4h_intersect, sna_2-4h_intersect, twi_2-4h_Toll^10b^ and sna_2-4h_Toll^10b^ additionally included initial germ band elongation (stage seven, 180-190 minutes) (Campos-Ortega and Hartenstein, 1997; Sandmann et al., 2007; Zeitlinger et al., 2007); twi_2-4h_Toll^10b^ and sna_2-4h_Toll^10b^ may have also included stages eight (190-220 minutes) and nine (220-260 minutes) (Campos-Ortega and Hartenstein, 1997; Zeitlinger et al., 2007). Twi_2-4h_intersect and sna_2-4h_intersect were tightly staged between stages 5-7 (Sandmann et al., 2007). Additional to stages four, five and six, twi_1-3h_hiConf may have included the latter part of stage two (preblastoderm, 25-65 minutes) and stage three (pole bud formation, 65-80 minutes) (Campos-Ortega and Hartenstein, 1997). The twi_4-6h_intersect dataset was restricted to stages eight to nine which included germ band elongation and segmentation of neuroblasts (Campos-Ortega and Hartenstein, 1997; Sandmann et al., 2007). Therefore, there were differences in the biological material used across TF_ALL.

The Notch signalling modifiers analysed in this study were selected based on identification in at least two of the screens reported in (Guruharsha et al., 2012).

### Breast cancer transcriptome datasets and molecular subtypes

Primary breast tumour gene expression data were downloaded from NCBI GEO (GSE12276, GSE21653, GSE3744, GSE5460, GSE2109, GSE1561, GSE17907, GSE2990, GSE7390, GSE11121, GSE16716, GSE2034, GSE1456, GSE6532, GSE3494, GSE68892 (formerly geral-00143 from caBIG)). All datasets were Affymetrix U133A/plus 2 chips and were summarised with Ensembl alternative CDF (Dai et al., 2005). RMA normalisation (Irizarry et al., 2003) and ComBat batch correction (Johnson et al., 2007) were applied to remove dataset-specific bias as previously described (Moleirinho et al., 2013; Sims et al., 2008). Intrinsic molecular subtypes were assigned based upon the highest correlation to Sorlie centroids (Sørlie et al., 2003), applied to each dataset separately. Centred average linkage clustering was performed using the Cluster and TreeView programs (Eisen et al., 1998). Centroids were calculated for each gene based upon the mean expression across each of the Sorlie intrinsic subtypes (Sørlie et al., 2003). These expression values were squared to consider up and down regulated genes in a single analysis. Orthology to the DroFN network was defined using Inparanoid (Östlund et al., 2009). Differential expression was calculated by t-test comparing normalised (unsquared) expression values in normal-like and basal-like tumours with false discovery rate correction (Benjamini and Hochberg, 1995).

### Invasion assays for validation of genes selected from NetNC results

MCF-7 Tet-On cells were purchased from Clontech and maintained as previously described (Liu et al., 2013).To analyse the ability of transfected MCF7 breast cancer cells to degrade and invade surrounding extracellular matrix, we performed an invasion assay using the CytoSelect™ 24-Well Cell Adhesion Assay kit. This transwell invasion assay allow the cells to invade through a matrigel barrier utilising basement membrane-coated inserts according to the manufacturer’s protocol. Briefly, MCF7 cells transfected with the constructs (Doxycycline-inducible *SNAI1* cDNA or *SNAI1 shRNA* with or without candidate gene cDNA) were suspended in serum-free medium. *SNAI1* cDNA or *SNAI1 shRNA* were cloned in our doxycyline-inducible pGoldiLox plasmid (pGoldilox-Tet-ON for cDNA and pGolidlox-tTS for shRNA expression) using validated shRNAs against *SNAI1* (NM_005985 at position 150 of the transcript (Liu et al., 2013)). pGoldilox has been used previously to induce and knock down the expression of *Ets* genes (Peluso et al., 2017).

Following overnight incubation, the cells were seeded at 3.0×10^5^ cells/well in the upper chamber and incubated with medium containing serum with or without doxycyline in the lower chamber for 48 hours. Concurrently, 10^6^ cells were treated in the same manner and grown in a six well plate to confirm over-expression and knockdown. mRNA was extracted from these cells and quantitative real-time PCR (RT-qPCR) was performed as previously described (Essafi et al., 2011); please see Supplemental File 2 for gene primers. The transwell invasion assay evaluated the ratio of CyQuant dye signal at 480/520 nm in a plate reader of cells from the two wells and therefore controlled for potential proliferation effects associated with ectopic expression. We used empty vector (mCherry) and scrambled shRNA as controls and to control for the non-specific signal. At least three experimental replicates were performed for each reading.

## ACKNOWLEDGEMENTS

IMO is grateful to Prof Jeremy Gunawardena and Prof Peter Sorger for hosting his visit to HMS and for helpful discussions. Thanks to Prof PS Thiagarajan, Prof Andrew Millar, Prof Wendy Bickmore, Prof Nick Hastie, Prof Ben Lehner and Prof Julian Dow for invaluable comments. Mr Nick Moir and Dr Seanna McTaggart assisted with testing the NetNC software distribution. We are grateful for funding from Medical Research Council (MC_UU_12018/25; IMO), Royal Society of Edinburgh Scottish Government Fellowship cofunded by Marie Curie Actions (IMO), Marie Curie Fellowship (BH), Breast Cancer Now (AHS). AE was supported by a Wellcome Trust Beit Memorial Fellowship (AE) and by funding from Prof. Nick Hastie’s laboratory (MC_PC_U127527180).

## AUTHOR CONTRIBUTIONS

IMO conceived the overall project, obtained funding, designed the computational and statistical aspects, implemented and benchmarked the NetNC algorithm, performed analysis of all TF datasets and invasion assay data, interpreted results, produced Figures 1, 3, 4, 6, produced all Tables except as noted below, performed orthology mapping, annotated the heatmap features in Figure 5 and supervised JO, BH, ALRL, MJF, EP-C. JO implemented the iterative minimum cut, co-designed and implemented the NetNC parameter optimisation, assisted with NetNC benchmarking and produced Figures 2, S9, Table S8. BH obtained funding, co-designed and implemented the DroFN network inference, benchmarking and produced Figure S1. MJF co-designed and implemented the comparison of NetNC against the MCL algorithm, produced Table S9. ALRL co-designed and implemented the Gaussian Mixture Modelling aspects of NetNC and co-designed Equation 9. IO, JO and ALRL wrote the NetNC software distribution. AHS obtained funding, co-designed and implemented the breast cancer transcriptome analysis, interpreted results, produced Figures 5 and S6. AE obtained funding, interpreted results, designed and performed all bench laboratory experiments including tissue culture, transfection and transwell assays. EP-C assisted with annotation, visualisation and interpretation of the NetNC-FTI networks, including production of Figure S5. IO led the writing of the manuscript and revised it for important intellectual content with input from JO, AHS, AE, BH, EP-C, ALRL.

## Supplemental Material for Overton *et al.* ‘Coherent Transcription Factor Target Networks Illuminate Epithelial Remodelling’

**Figure S1.**
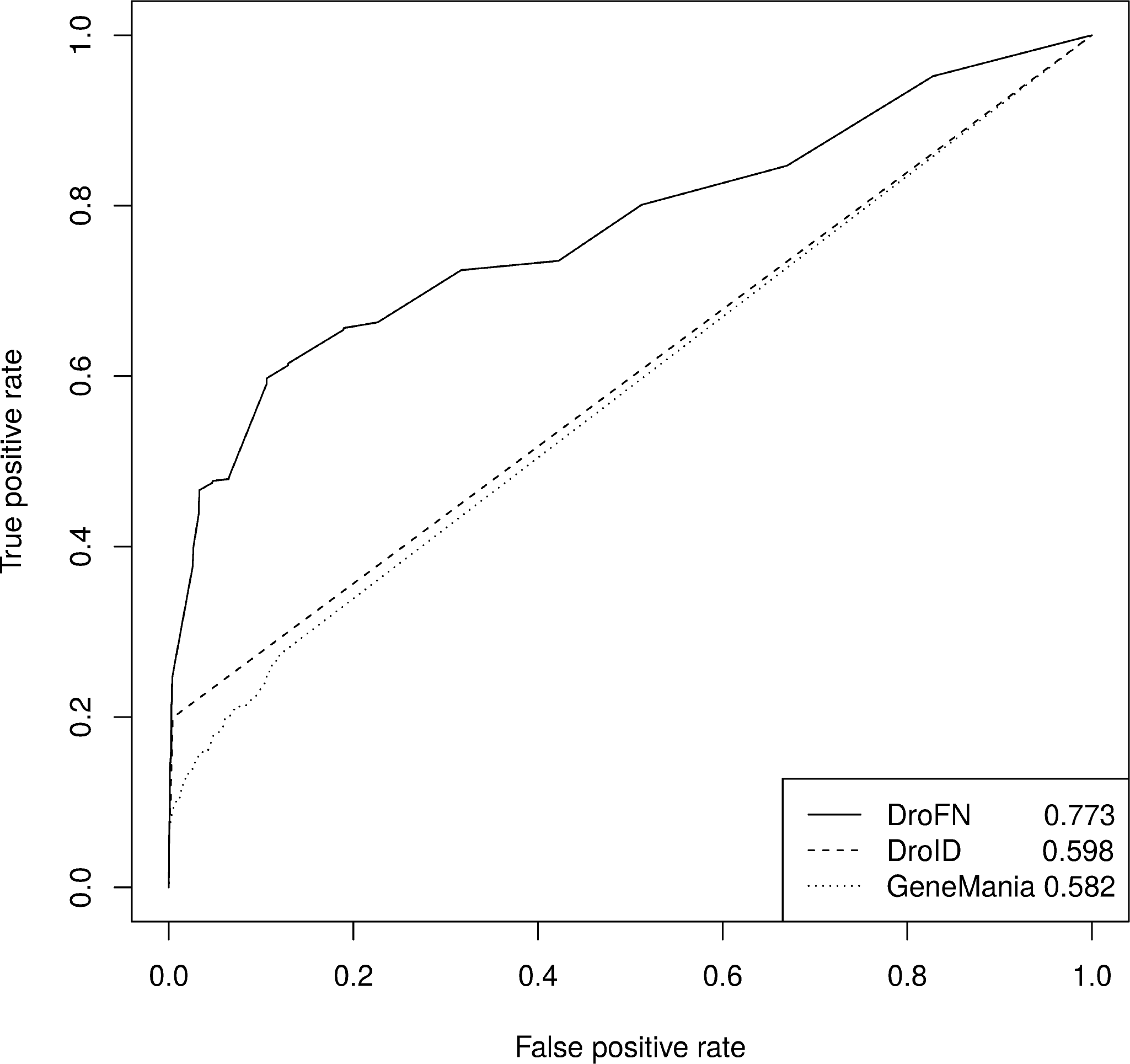
Related to Methods. Reciever Operator Characteristic (ROC) curves for Functional Gene Networks on time separated blind test data (TEST-NET). ROC curves for DroFN, DroID and GeneMania networks are shown with respective AUC values of 0.773, 0.598 and 0.582. DroFN has significantly larger AUC than DroID (*p*<2.13×10^−11^) and GeneMania (*p*<3.2×10^−22^) on TEST-NET.

**Figure S2.**
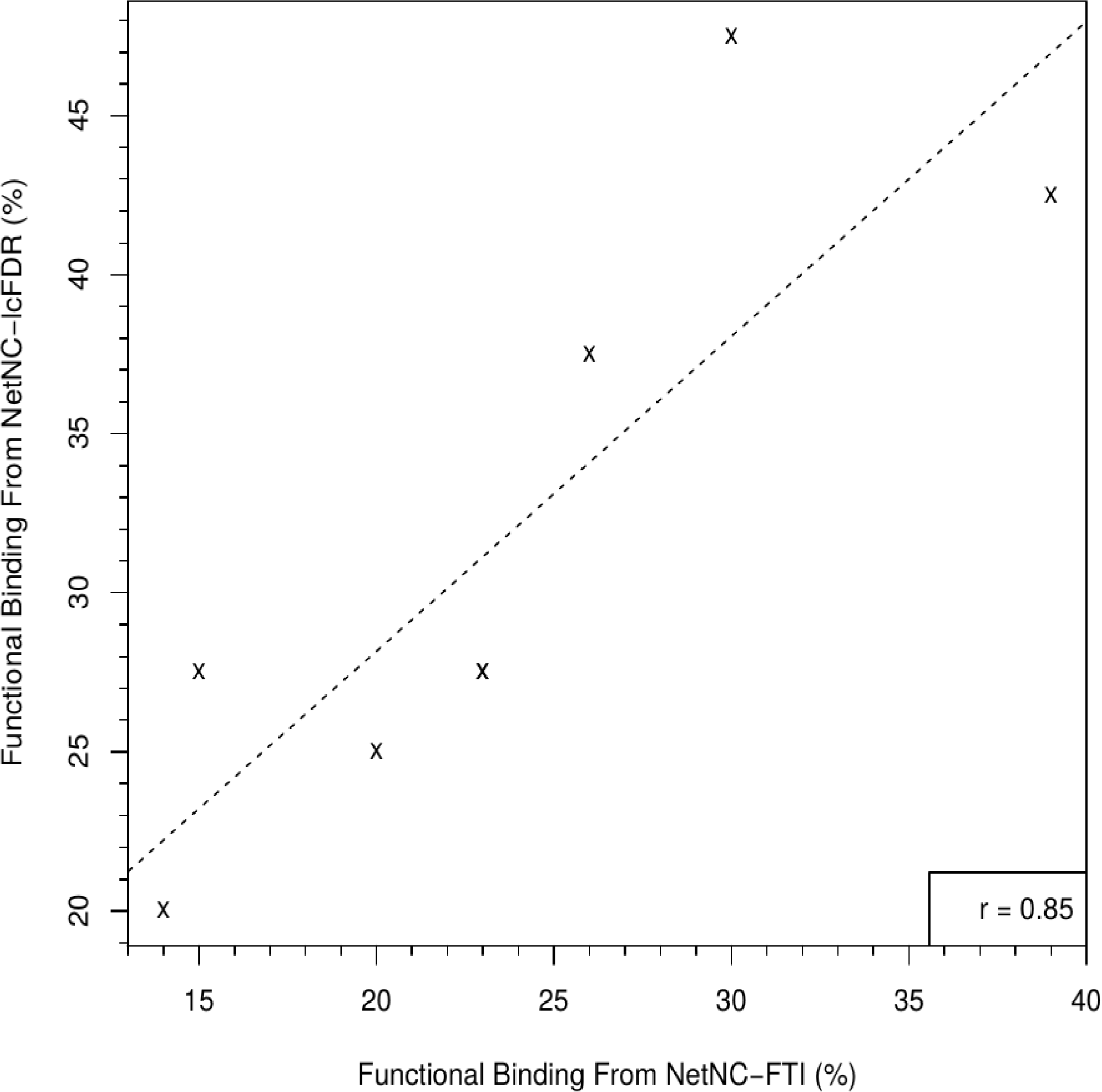
Related to Table 1. Agreement between NetNC-lcFDR and NetNC-FTI estimates of functional binding. The percentage of predicted functional binding from NetNC-lcFDR and NetNC-FTI for each of the nine TF_ALL datasets is shown as crosses (‘X’). Results from the two different methods correlated well (r=0.9, *p=*0.008) and had median difference of only 5.5%. This concordance supports the results from both NetNC-FTI and NetNC-lcFDR.

**Figure S3.**
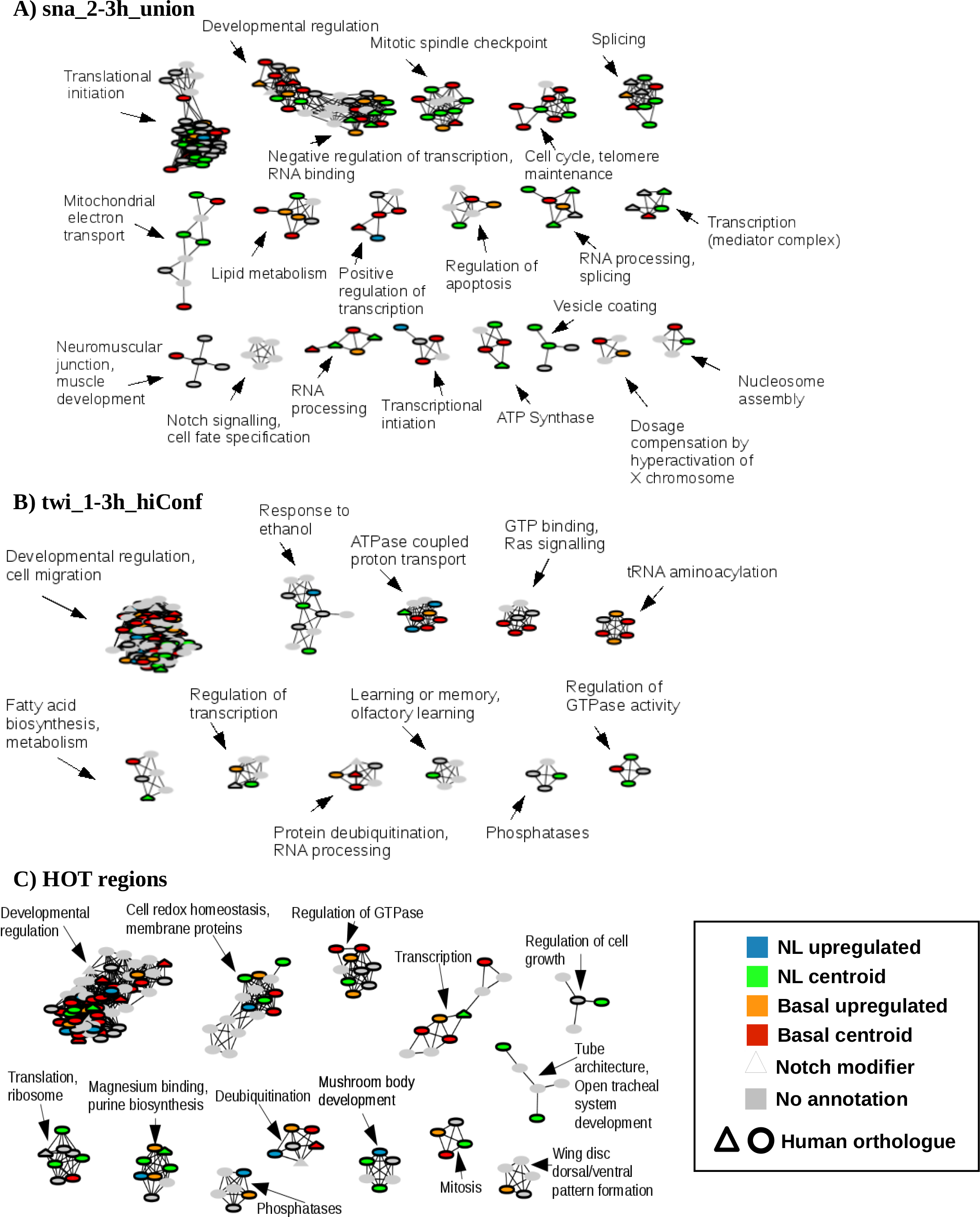
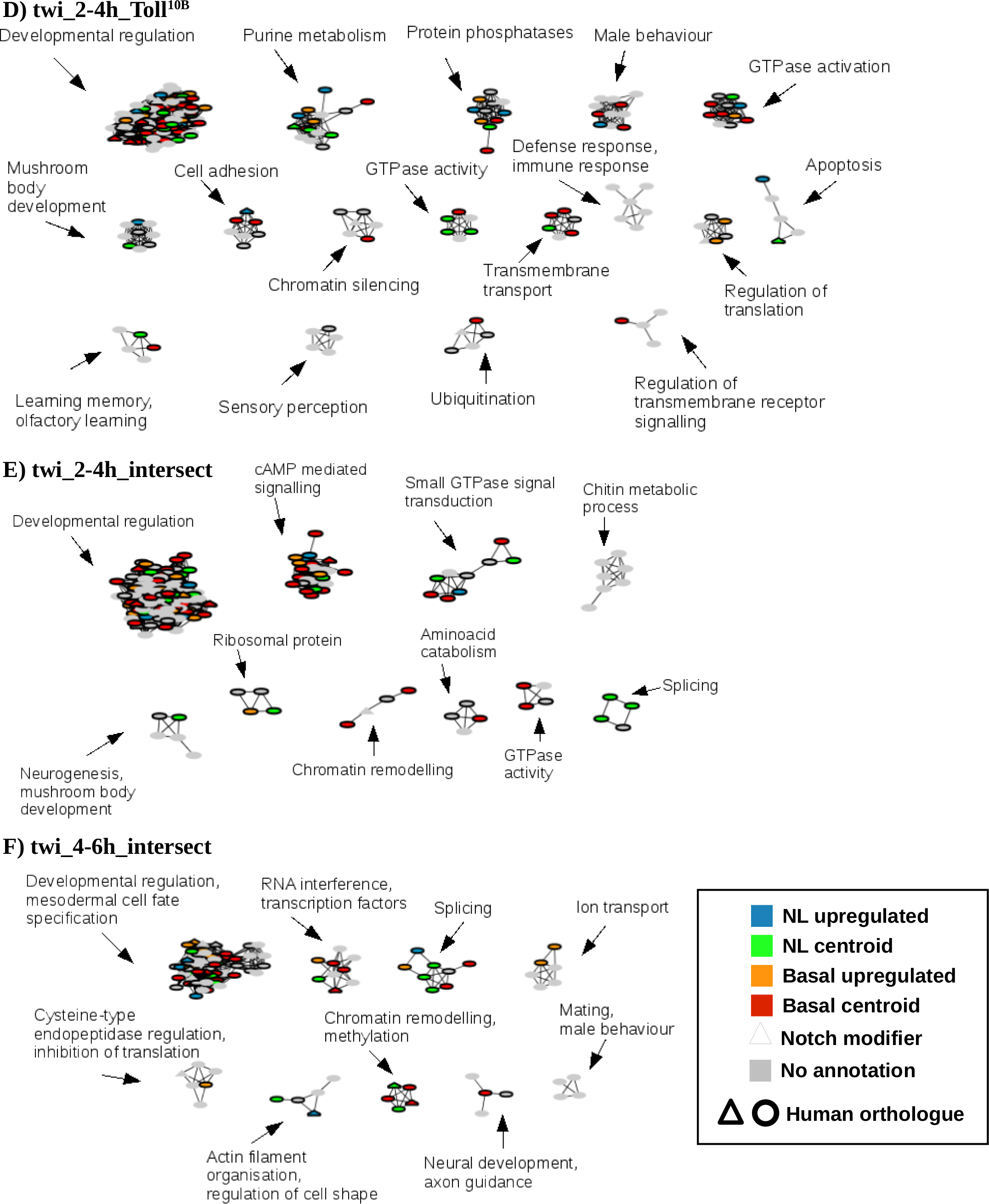
Related to Figure 4 and Supplemental File 1. Networks of functionally coherent candidate target genes identified by NetNC-FTI. Networks of functionally coherent candidate target genes identified by NetNC-FTI. The key (bottom right) indicates annotations for human orthology (bold node border) and Notch screen hits (triangular nodes). Many orthologues were assigned to either basal-like (BL, red) or normal-like centroids (NL, green); otherwise, node colour indicates upregulated gene expression in NL (blue) compared to BL (orange) subtypes (*q*<0.05) or no annotation to the BL or NL subtypes (grey). Clusters with at least four members are shown, cytoscape sessions with full NetNC-FTI results are given in Additional File 4. Very few clusters were composed entirely from genes identified only in a single dataset, examples included: snoRNAs/nucleolar proteins (twi_2-3h_union), transferases (HOT), defense response/immune response (twi_2-4h_Toll^10b^) and chitin metabolism (twi_2-4h_intersect).

**Figure S4.**
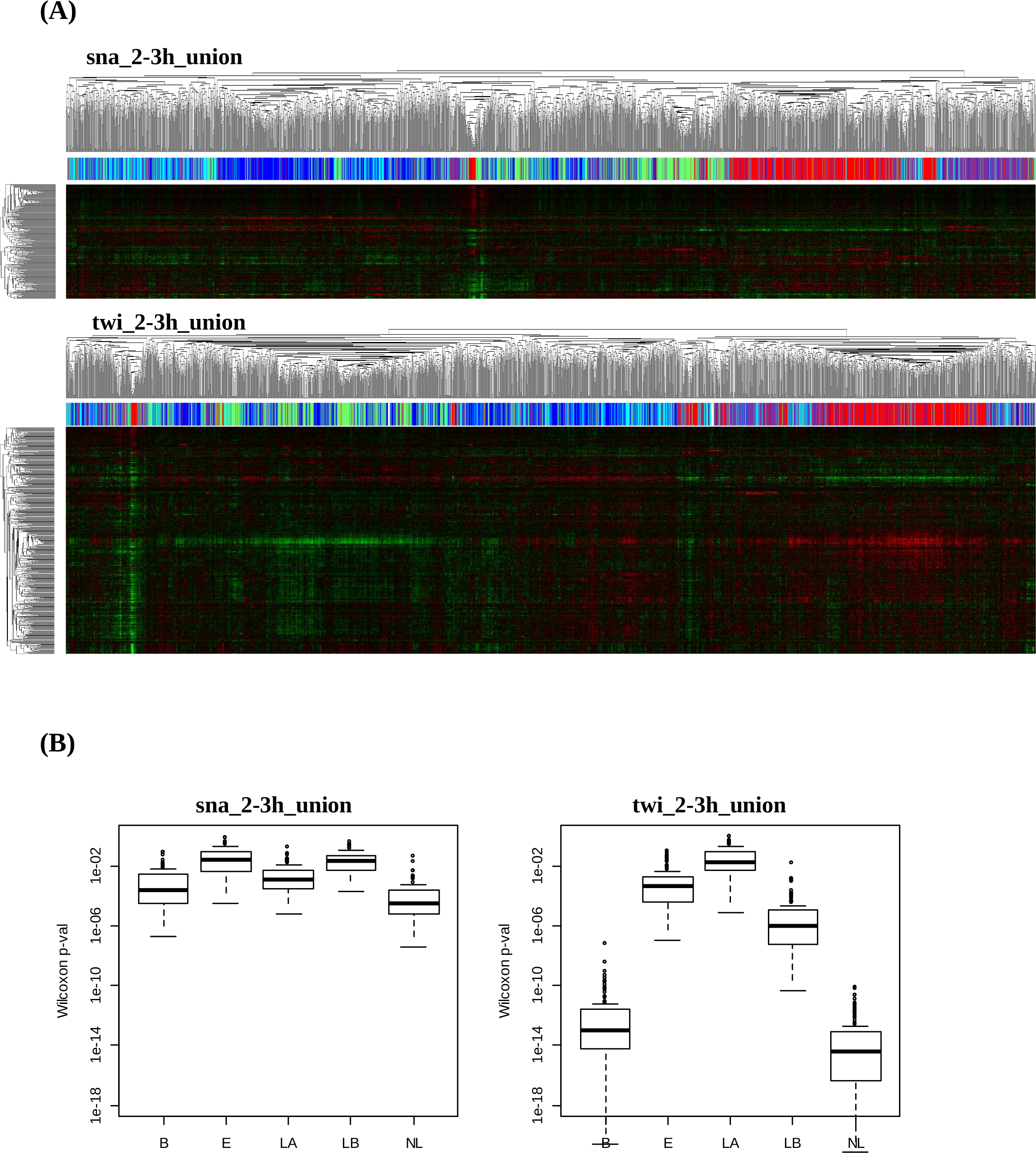
Related to Figure 5. Unsupervised clustering of 2999 primary breast cancers with NetNC results for individual Snail and Twist datasets. (A) Data shown represent results clustering of primary breast tumour gene expression data (detailed in Methods, section 4.4) with orthologous genes from NetNC-FBT analysis of sna_2-3h_union (top, n=256) and twi_2-3h_union (bottom, n=467). Intrinsic subtypes are shown along the top of each heatmap, colours correspond to: HER2-overexpressing (purple), basal-like (red), normal-like (green), luminal A (dark blue) and luminal B (light blue). Clustering based on sna_2-3h_union shows better separation of clusters than twi_2-3h_union. (B) Significantly higher squared centroid values for multiple breast cancer subtypes (Wilcoxon *p*<0.05) were obtained when compared with resampled orthologues for NetNC-FBT analysis of sna_2-3h_union (left, n=256) and twi_2-3h_union (right, n=467); 100 resamples were performed. Breast cancer subtypes are indicated on the x-axis of each plot as follows: B (basal-like), E (HER2-overexpressing), LA (luminal A), LB (luminal B), NL (normal-like). Similar patterns were observed between these two datasets, with strongest enrichment for basal-like and normal-like centroids; sna_2-3h_union also showed relatively stronger enrichment for the luminal A centroid compared with results for twi_2-3h_union. Absolute differences in *p* between sna_2-3h_union and twi_2-3h_union may reflect the difference in dataset sizes (n=256 vs n=467), indeed stronger p-values are typically associated with comparisions across larger datasets.

**Figure S5.**
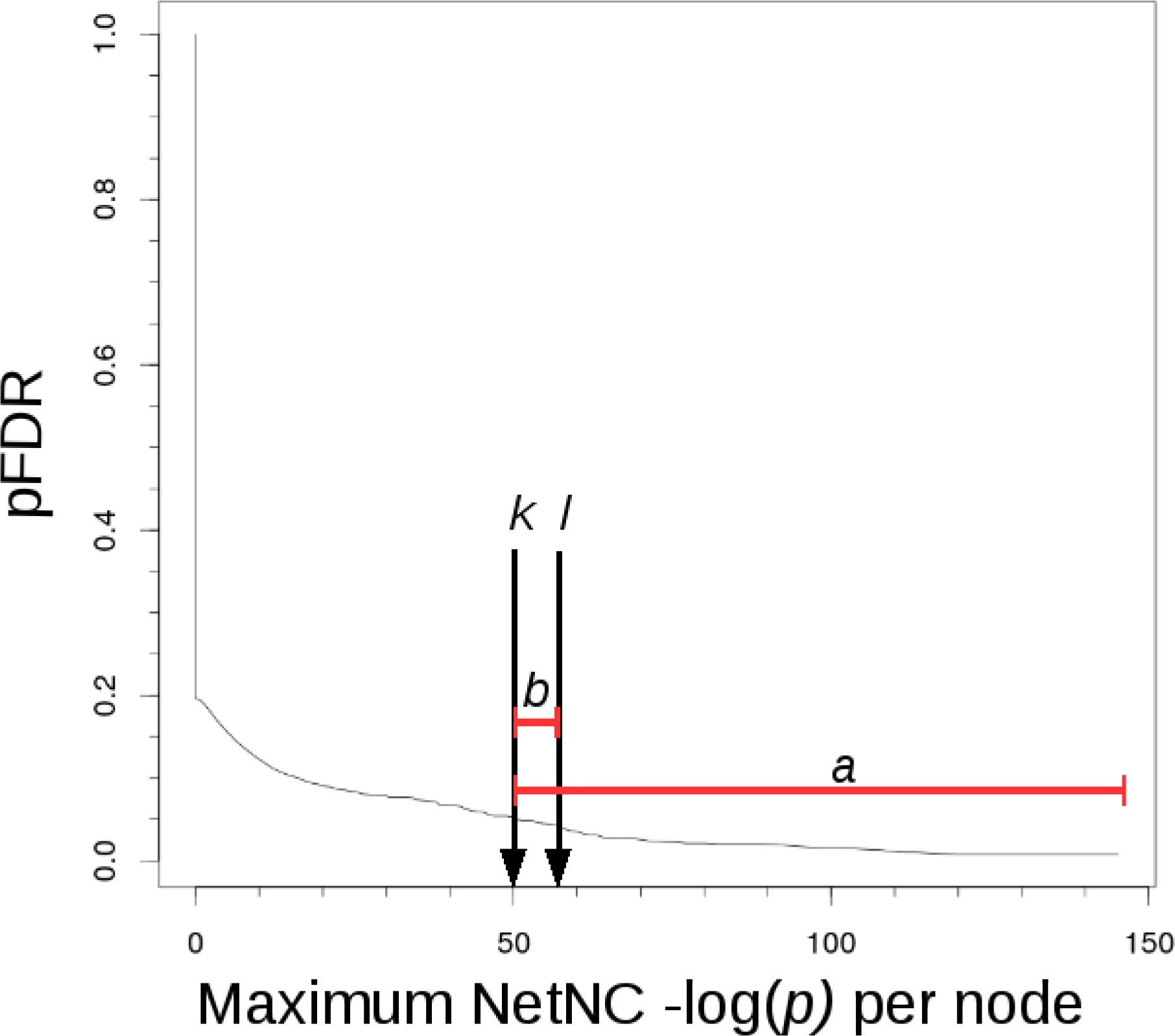
Related to Methods. Estimation of local False Discovery Rate (lcFDR) from positive False Discovery Rate. To estimate lcFDR the expected number of false positives (FP) is first estimated for the edges (*X*) under *b* (red bounding line). The pFDR at *k* (pFDR_*k*_) multiplied by the number of edges (*n*) under *a* (red bounding line) gives the FP above threshold *k* (FP_*k*_); the pFDR at *l* (pFDR_*l*_) multiplied by (*n – X*) gives the expected number of false positives for threshold *j* (FP_*l*_). Therefore, lcFDR bounded by *k* and *l* is estimated as (FP_*k*_ – FP_*l*_) / *X.* We took values of *k* and *l* such that the interval (*k, l*) contained only edges that had *p*≡*k.* Values for the twi_2-6h_intersect dataset are shown.

**Figure S6.**
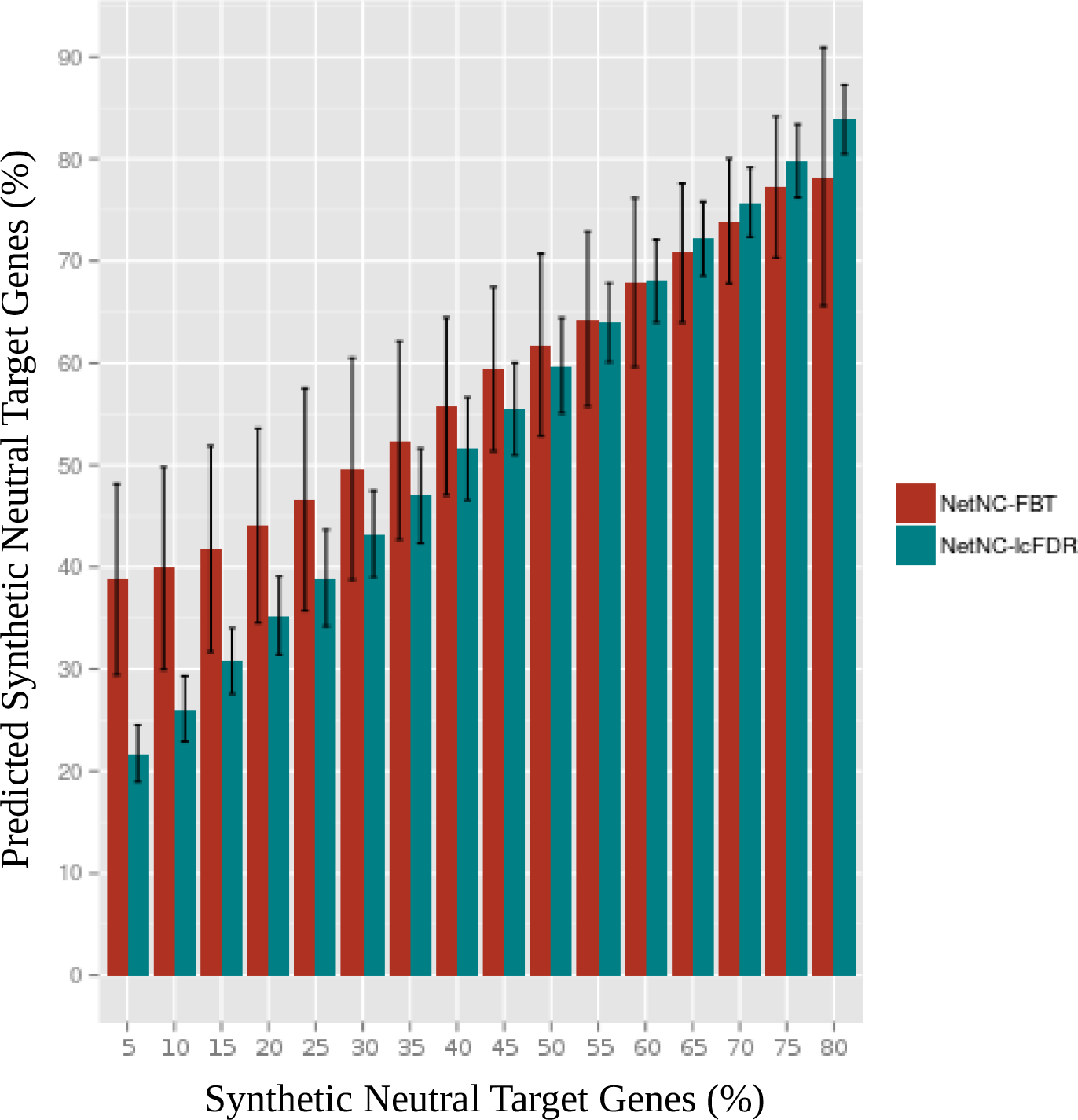
Related to Methods. Proportion of neutral binding predicted by NetNC-FBT and NetNC-lcFDR. The proportion of Synthetic Neutral Target Genes (SNTGs) in the TEST-CL_8PW datasets analysed is shown on the x-axis. The predicted proportion of SNTGs by NetNC-FBT (red) and NetNC-lcFDR (blue-green) is shown on the y-axis. Error bars represent 95% confidence intervals calculated over 100 resamples. Systematic overestimation of %SNTG was greatest for the NetNC-FBT approach which also had large 95% confidence intervals relative to NetNC-lcFDR.

**Figure S7.**
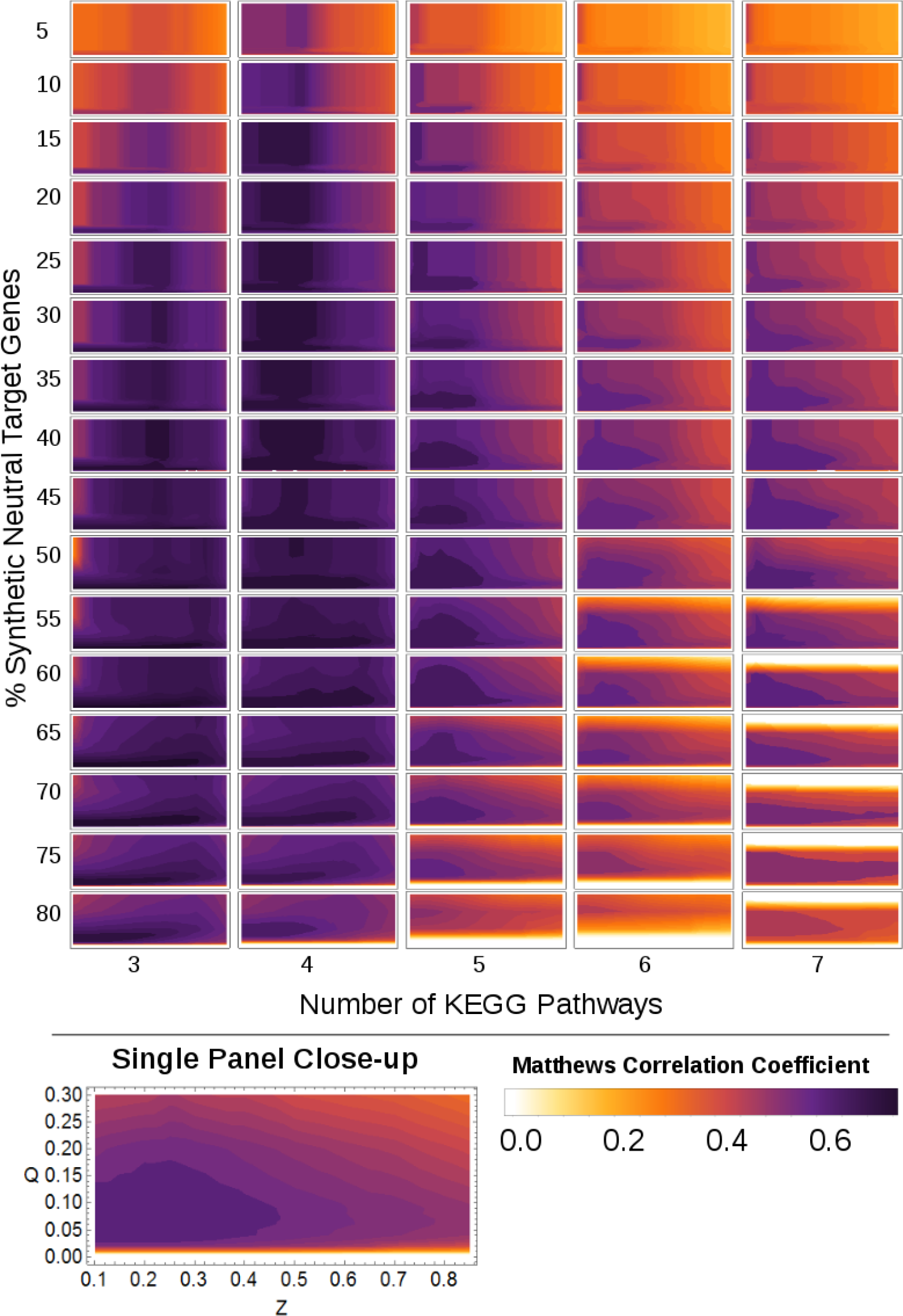
Related to Methods. NetNC Functional Target Identification Performance on TRAIN-CL_ALL. A range of values for the number of KEGG pathways (columns) and %Synthetic Neutral Target Genes (rows) were examined. Each invividual panel shows NetNC-FTI performance across values of the Q (FDR) and Z (graph density) parameters. Contours show the mean Matthews Correlation Coefficient value for the 100 resamples per individual panel. The NetNC-FTI parameters (Q, Z), were optimised on these data as described in the main manuscript Methods section 4.2.3.

**Table S1.** Related to Methods. Evaluation of DroFN on Time Separated Blind Test Data (TEST-NET). Column headings: Matthews correlation coefficient (MCC), false positive rate (FPR), true positive rate (TPR), area under the Receiver Operator Characteristic curve (AUC). DroFN performed best on the data examined and had FPR close to the functional interaction prior estimated from the training data (0.044). Values of AUC for DroFN were significantly better than DROID (*p=*2.13×10^−11^) or GeneMania (*p=*3.19×10^−22^).

| Network | MCC | FPR | TPR | AUC |
| --- | --- | --- | --- | --- |
| DroFN | 0.448 | 0.047 | 0.475 | 0.773 |
| DROID | 0.383 (max 0.385) | 0.0046 | 0.199 | 0.598 |
| GeneMania | 0.133 (max 0.243) | 0.121 | 0.274 | 0.582 |

**Table S2.**
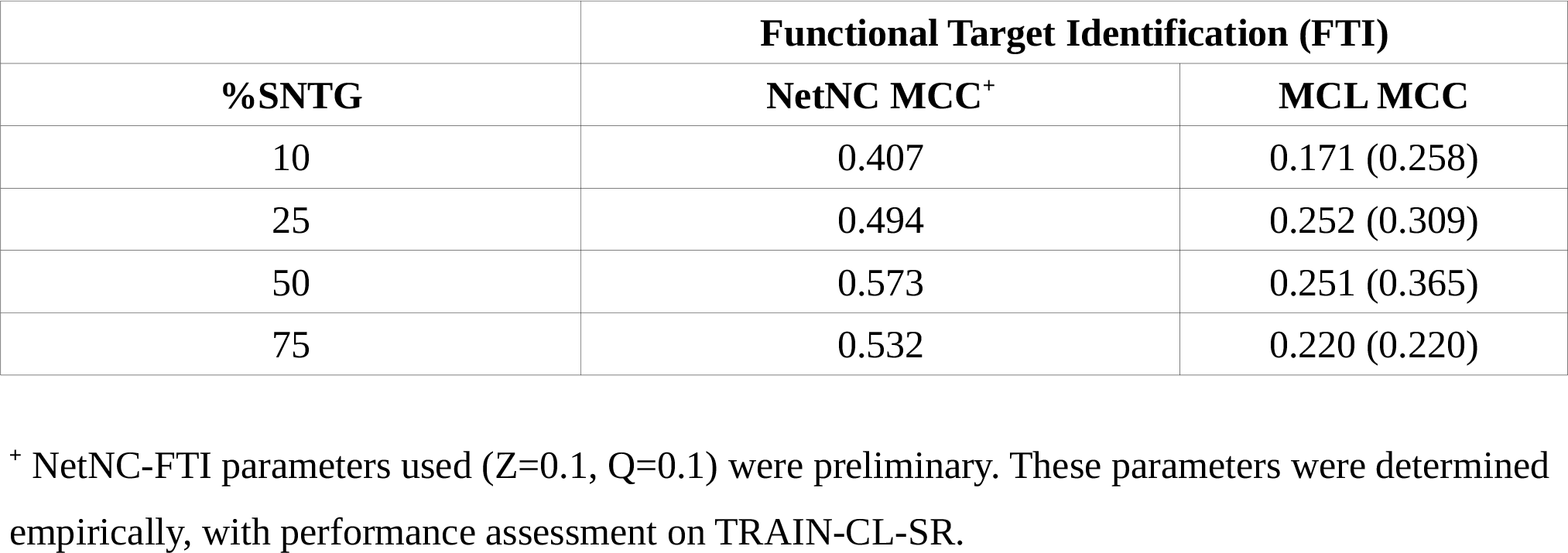
Related to Figure 2. Results from preliminary study of NetNC and MCL performance on TEST-CL-SR. MCC, Matthews Correlation Coefficient. MCL performance for overall best-performing inflation value (I=3.6) is given at FDR *p*<0.05. Performance values in brackets represent taking inflation values that were optimal at the value of %SNTG on TRAIN-CL-SR. Results represent performance on TEST-CL-SR, which comprised a single resample at each %SNTG value examined. MCC values for NetNC performance were considerably higher than those obtained using the MCL algorithm.

**Table S3.**
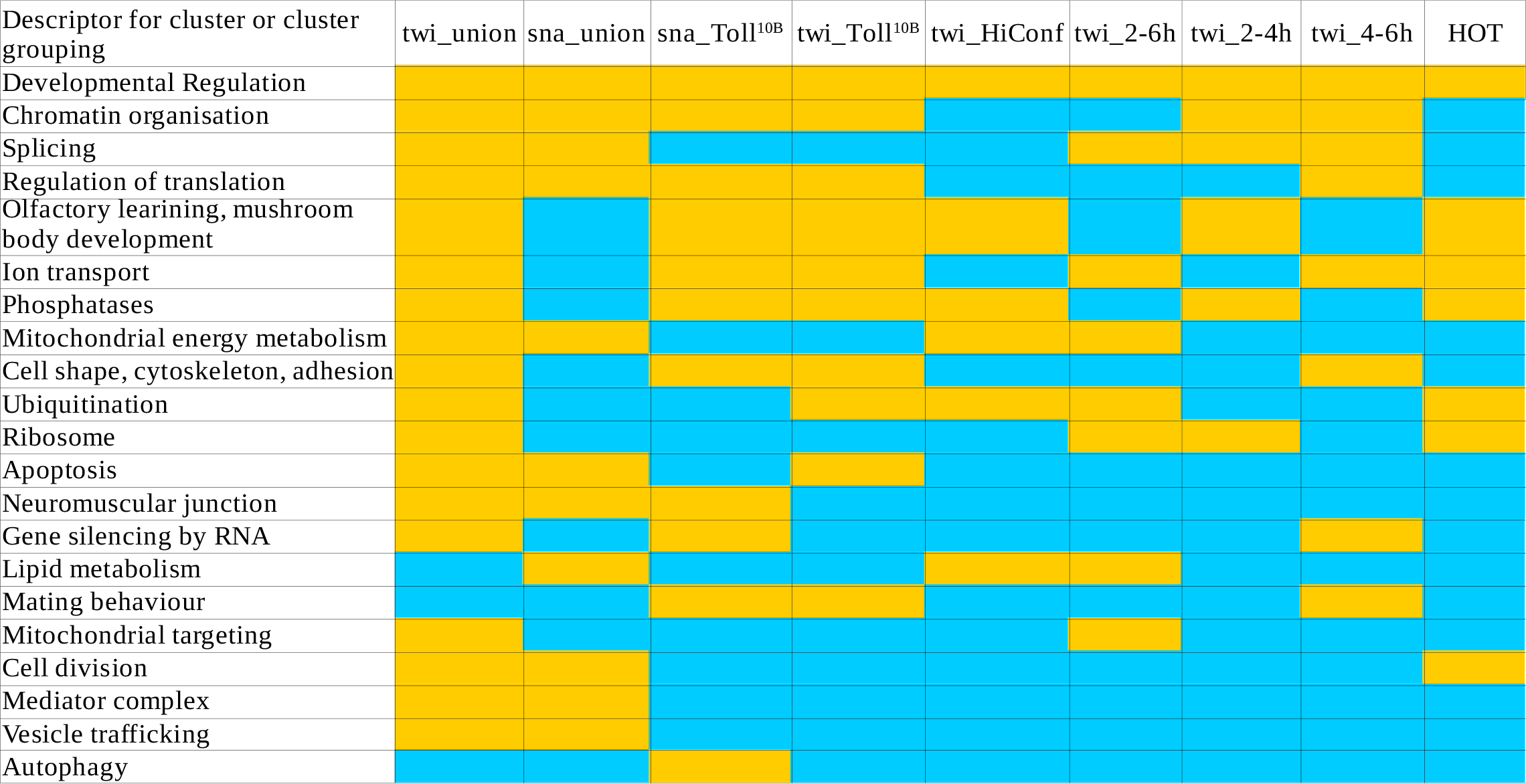
Related to Figure 4 and Figure S3. Overall summary of clusters identified by NetNC-FTI across the nine TF_ALL datasets. Clusters and cluster groupings are listed on the left of the table, orange table cells indicates that the biological grouping was found in NetNC-FTI results for the corresponding dataset, blue table cells indicate that the grouping was not identified.

**Table S4.**
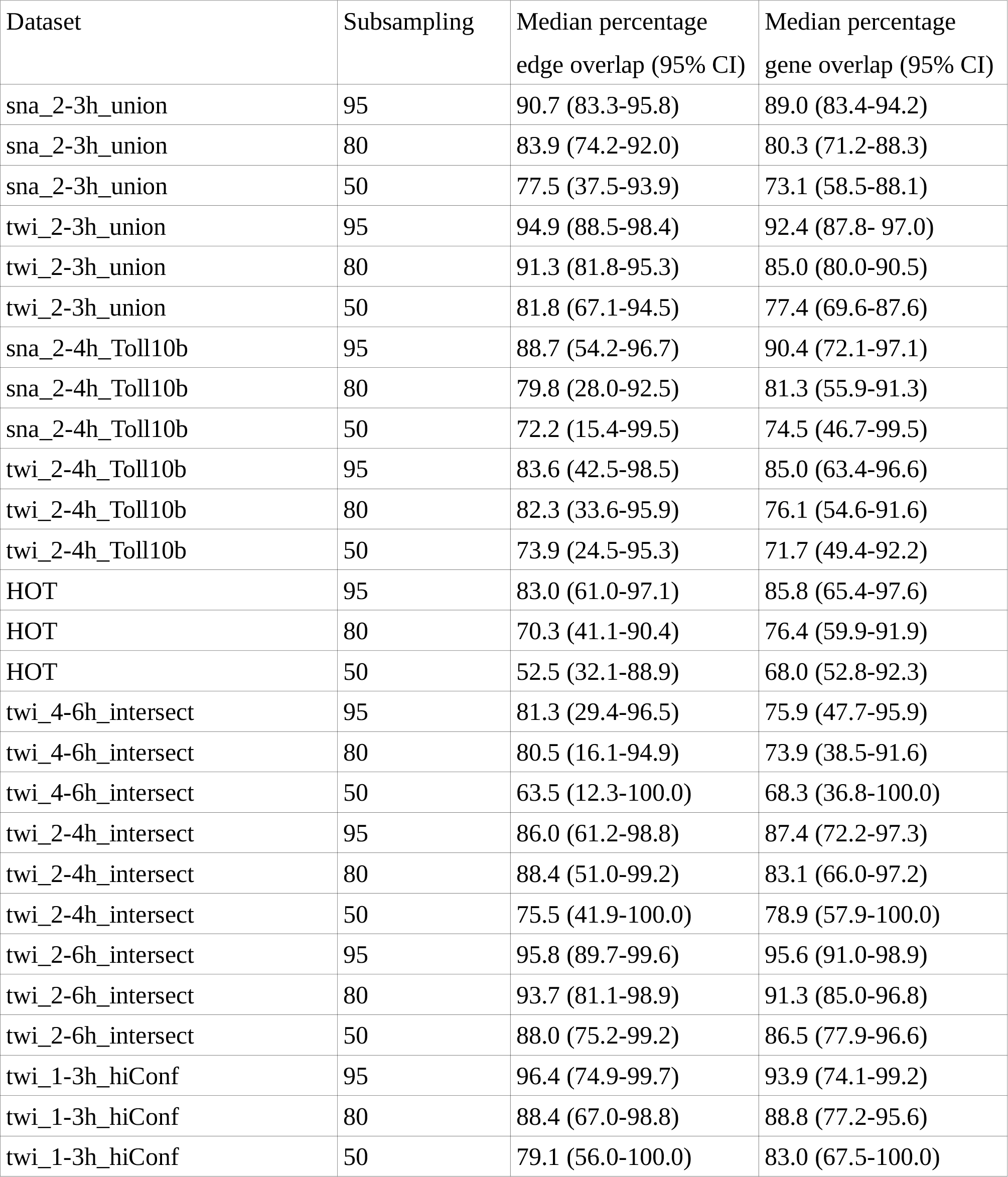
Related to Figure 4 and Figure S3. Overlap for NetNC-FTI subsamples of the TF_ALL datasets. Data represents the proportion of edges and genes returned by NetNC-FTI for the subsampled dataset that were also present in NetNC-FTI results for the full dataset (i.e. at 100%). At subsampling rates 95%, 80%, 50%, the average edge overlap values across TF_ALL were respectively 91%, 84% and 77%; derived as the median value (average) of the median overlap per dataset (respective median 95% CI 83-96%, 74-94%, 37-92%). At subsampling rates 95%, 80%, 50%, the average overlap values across TF_ALL were respectively 89%, 81% and 75%; derived as the median value (average) of the median overlap per dataset. (respective median 95% CI 72%-97.2%, 66%-92%, 58%-97%). Some subsamples taken as input to NetNC had low overlap with the NetNC-FTI reference output (reference_net) for any given complete input dataset. Indeed, the reference_net represented between 14% to 39% of the total input gene list across the nine TF_ALL datasets. Subsamples that excluded a high proportion of the nodes in reference_net would be expected to result in weaker hypergeometric mutual clustering values for the remaining nodes found in reference_net because these nodes would be expected to have relatively few common neighbours. Therefore, subsampling of the input gene list is expected to produce NetNC results that have reduced overlap with reference_net; this effect is also a source of variation in overlap across subsamples, reflected in the 95% CI values. Also, the probability of sampling nodes in reference_net is lower when a smaller fraction of the complete input TF_ALL gene list is covered by reference_net, leading to a greater subsampling-associated loss of nodes and edges. Consistent with this interpretation, TF_ALL datasets with the highest NetNC-FTI functional binding proportion (twi_1-3h_hiConf, twi_2-6h_intersect, HOT; see Table 1) were less sensitive to subsampling than datasets with relatively low predicted functional binding such as sna_2-4h_Toll^10b^ and twi_4-6h_intersect.

**Table S5.** Related to Figure 4 and Figure S3. Frequently identified genes within the Developmental Regulation Clusters. The thirteen genes in this table (DRC-13) were found in the developmental regulation cluster returned by NetNC-FTI in least seven of the nine TF_ALL datasets.

| Identifier | Name (count in TF_ALL) | Uniprot Function when available, otherwise the Flybase Summary is given (indicated by *) |
| --- | --- | --- |
| FBgn0004009<br>Current ID:<br>FBgn0284084 | Wingless (9/9) | Binds as a ligand to a family of frizzled seven-transmembrane receptors and acts through a cascade of genes on the nucleus.<br>Segment polarity protein. May be a growth factor. Acts on neighboring cells to regulate at least one gene, the homeobox segmentation gene engrailed. Wg signal represses arm phosphorylation. Wg signaling operates by inactivating the sgg repression of engrailed autoactivation. Wg and Wnt2 have a role in the developing trachea and together are responsible for all dorsal trunk formation. Wg also acts in the developing epidermis. Acts as a morphogen, and diffuses long distances despite its lipidation. Lipophorin is required for diffusion, probably by acting as vehicle for its movement, explaining how it can spread over long distances despite its lipidation. In non-neuronal cells, wls directs wg secretion via clathrin-mediated endocytosis and the retromer complex (a conserved protein complex consisting of Vps26 and Vps35) to sustain a wls traffic loop encompassing the Golgi, the cell surface, an endocytic compartment and a retrograde route leading back to the Golgi. In neuronal cells (the larval motoneuron NMJ), wg signal moves across the synapse through the release of wls-containing exosome-like vesicles. |
| FBgn0004647 | Notch (8/9) | Signaling protein, which regulates, with both positive and negative signals, the differentiation of at least central and peripheral nervous system and eye, wing disk, oogenesis, segmental appendages such as antennae and legs, and muscles, through lateral inhibition or induction. Functions as a receptor for membrane-bound ligands Delta and Serrate to regulate cell-fate determination. Upon ligand activation, and releasing from the cell membrane, the Notch intracellular domain (NICD) forms a transcriptional activator complex with Su(H) (Suppressor of hairless) and activates genes of the E(spl) complex. Essential for proper differentiation of ectoderm. Fringe (fng) acts in the Golgi to determine the type of O-linked fucose on the EGF modules in N, altering the ability of N to bind |
|  |  | with Delta (Dl). O-fut1 also has a role in modulating the interaction. Rumi acts in the endoplasmic reticulum to glucosylate the EGF modules in N, this is required for correct folding and cleavage of N. |
| FBgn0000659 | fork head<br>(8/9) | Fkh promotes terminal as opposed to segmental development. In the absence of fkh, this developmental switch does not occur. The nuclear localization of the fkh protein suggest that fkh regulates the transcription of other, subordinate, genes. |
| FBgn0010389 | heartless<br>(7/9) | May be required for patterning of muscle precursor cells. May be essential for generation of mesodermal and endodermal layers, invaginations of various types of cells and CNS formation. |
| FBgn0011225 | jaguar (7/9) | Myosin is a protein that binds to actin and has ATPase activity that is activated by actin. Together CLIP-190 and jar may coordinate the interaction between the actin and microtubule cytoskeleton. May link endocytic vesicles to microtubules and may be involved in transport in the early embryo and in the dynamic process of dorsal closure. It is believed that its function changes during the life cycle. |
| FBgn0000577 | engrailed<br>(7/9) | This protein specifies the body segmentation pattern. It is required for the development of the central nervous system. Transcriptional regulator that represses activated promoters. Wg signaling operates by inactivating the SGG repression of EN autoactivation. |
| FBgn0003118 | pointed (7/9) | Required for glial-neuronal cell interactions at the ventral midline which are necessary for the proper elaboration of commissures in the embryonic CNS. |
| FBgn0017397<br>Current ID:<br>FBgn0264491 | held out<br>wings (7/9) | Required for integrin-mediated cell-adhesion in wing blade. Vital role in steroid regulation of muscle development and to control heart rate. Required during embryogenesis, in late stages of somatic muscle development, for myotube migration and during metamorphosis for muscle reorganization. |
| FBgn0003254 | ribbon (7/9) | *Ribbon is a nuclear BTB-domain protein, expressed in most embryonic cells. It is required for development of the salivary gland and trachea, as well as for dorsal closure. It regulates both growth and differentiation of salivary gland cells. |
| FBgn0000014 | abdominal A<br>(7/9) | Sequence-specific transcription factor which is part of a developmental regulatory system that provides cells with specific |
|  |  | positional identities on the anterior-posterior axis. Required for segmental identity of the second through eighth abdominal segments. Once a pattern of abd-A expression is turned on in a given parasegment, it remains on the more posterior parasegment, so that the complex pattern of expression is built up in the successive parasegments. Appears to repress expression of Ubx whenever they appear in the same cell, but abd-A is repressed by Abd-B only in the eight and ninth abdominal segments. |
| FBgn0003900 | twist (7/9) | Involved in the establishment and dorsoventral patterning of germ layers in the embryo. |
| FBgn0003177<br>Current ID:<br>FBgn0262614 | polychaetoid (7/9) | *Polychaetoid is a broadly acting protein that is associated with multiple proteins at the surface and within the cytoskeleton, connecting events between the two. |
| FBgn0015513 | myoblast city (7/9) | *Myoblast city is an unconventional bipartite GEF that, in combination with Ced-12, activates the Rac1 monomeric GTPase. It contributes to myoblast fusion, border cell migration and dorsal closure. |

**Table S6.** Related to Figure 4 and Figure S3. NetNC functionally coherent TF targets found in chromatin organisation clusters across multiple datasets. The genes listed above were identified in NetNC-FTI chromatin organisation clusters for two or more of the TF_ALL datasets.

| Gene symbol | Identifier | Datasets where NetNC-FTI results include the given gene | Brief summary of function |
| --- | --- | --- | --- |
| *ph-d* | FBgn0004860 | sna_2-4h_Toll<sup>10b</sup>, twi_2-4h_Toll<sup>10b</sup>, twi_2-4h_intersect | Polycomb Repressive Complex 1 (PRC1) [Shao 1999] |
| *psc* | FBgn0005624 | sna_2-4h_Toll<sup>10b</sup>, twi_2-4h_Toll<sup>10b</sup>, twi_2-4h_intersect | Polycomb Repressive Complex 1 (PRC1) [Shao 1999] |
| *lolal* | FBgn0022238 | sna_2-4h_Toll<sup>10b</sup>, twi_2-4h_Toll<sup>10b</sup> | TrxG-related gene, regulates dorso-ventral patterning and is required for silencing at polycomb response elements [Mishra 2003, Quijano 2016] |
| *ph-p* | FBgn0004861 | sna_2-4h_Toll<sup>10b</sup>, twi_2-4h_Toll<sup>10b</sup> | Polycomb Repressive Complex 1 (PRC1) [Shao 1999] |
| *corto* | FBgn0010313 | sna_2-4h_Toll<sup>10b</sup>, twi_2-4h_Toll<sup>10b</sup> | physically interacts with PcG and TrxG proteins [Salvaing 2003, Lopez 2001] |
| *trithorax* | FBgn0003862 | twi_2-3h_union, twi_4-6h_intersect | histone methyltransferase (TrxG gene) [Tie 2014, Ingham and Whittle 1980] |
| *su(var)205* | FBgn0003607 | twi_2-3h_union, twi_4-6h_intersect | histone H3K9 binding, gene silencing factor [Fanti 1998, Fanti 2008] |
| *moira* | FBgn0002783 | twi_2-3h_union, twi_4-6h_intersect | TrxG gene involved in nucleosome organisation [Hong & Choi 2016, Crosby 1999] |
| *taranis* | FBgn0040071 | twi_2-3h_union, twi_4-6h_intersect | genetically interacts with TrxG and PcG genes [Schuster 2015, Calgaro 2002, Fauvarque 2001]. |
| *su(var)3-9* | FBgn0263755<br>(DroFN ID: FBgn0003600) | twi_2-4h_intersect, twi_2-4h_Toll<sup>10b</sup>, twi_2-3h_union | histone H3 methylase and gene silencing factor [Czermin 2001, Schotta 2002] |

**Table S7.** Related to Methods. Summary of pathways selected for gold standard data TRAIN-CL and TEST-CL.

| **Dataset** | **KEGG Pathway ID** | **Pathway Name** | **Number of Genes** |
| --- | --- | --- | --- |
| TRAIN-CL | dme03020 | RNA polymerase | 26 |
|  | dme03430 | Mismatch repair | 20 |
|  | dme04330 | Notch signalling | 22 |
|  | dme04914 | Progesterone-mediated oocyte maturation | 45 |
|  | dme00790 | Folate biosynthesis | 24 |
|  | dme00590 | Arachidonic acid metabolism | 15 |
|  | dme00280 | Valine, leucine and isoleucine degradation | 15 |
| TEST-CL | dme00510 | N-Glycan biosynthesis | 36 |
|  | dme00760 | Nicotinate and nicotinamide metabolism | 11 |
|  | dme03030 | DNA replication | 34 |
|  | dme04140 | Regulation of autophagy | 14 |
|  | dme04350 | TGF-beta signaling pathway | 29 |
|  | dme04745 | Phototransduction - fly | 27 |
|  | dme04122 | Sulfur relay system | 9 |
|  | dme00600 | Sphingolipid metabolism | 9 |

**Table S8.** Related to Methods. NetNC-FTI parameter values optimised for percentage Synthetic Neutral Target Genes (%SNTG). This table shows parameter values for NetNC-FTI optimised on TRAIN_ALL. These data may be used to specify NetNC-FTI parameters where an estimate of %SNTG is available, for example derived from NetNC-lcFDR analysis. In general there is a trend toward less stringent values of the optimised parameters (higher Q, lower Z) as the %SNTG increases.

| %SNTG | Optimised Z | Optimised Q |
| --- | --- | --- |
| 5 | 0.346 | 0.025 |
| 10 | 0.346 | 0.025 |
| 15 | 0.3735 | 0.0400171 |
| 20 | 0.3655 | 0.051006 |
| 25 | 0.3735 | 0.0612547 |
| 30 | 0.3615 | 0.0702808 |
| 35 | 0.3525 | 0.0834251 |
| 40 | 0.34 | 0.0920248 |
| 45 | 0.3325 | 0.0904339 |
| 50 | 0.3005 | 0.0681088 |
| 55 | 0.2945 | 0.0543454 |
| 60 | 0.25 | 0.0423662 |
| 65 | 0.2005 | 0.0546396 |
| 70 | 0.1355 | 0.0652494 |
| 75 | 0.2875 | 0.0991509 |
| 80 | 0.2535 | 0.2 |

**Table S9.** Related to Methods. Optimisation of MCL Inflation value for the Functional Target Identification task. Results shown represent performance on the TRAIN-CL-SR dataset. The overall best performing inflation value across all four TRAIN-CL datasets examined was 3.6.

| Inflation Value | Matthews Correlation Coefficient |  |  |  |
| --- | --- | --- | --- | --- |
|  | TRAIN-CL10% | TRAIN-CL25% | TRAIN-CL50% | TRAIN-CL75% |
| 2 | 0.303* | 0.318 | 0.376 | 0.271 |
| 2.2 | 0.241 | 0.348 | 0.269 | 0.253 |
| 2.4 | 0.217 | 0.357 | 0.418* | 0.270 |
| 2.6 | 0.186 | 0.367* | 0.402 | 0.359 |
| 2.8 | 0.174 | 0.340 | 0.400 | 0.365 |
| 3 | 0.161 | 0.300 | 0.341 | 0.343 |
| 3.2 | 0.201 | 0.323 | 0.370 | 0.355 |
| 3.4 | 0.204 | 0.355 | 0.384 | 0.368 |
| 3.6 | 0.196 | 0.344 | 0.406 | 0.393* |
| 3.8 | 0.196 | 0.333 | 0.392 | 0.390 |
| 4 | 0.193 | 0.330 | 0.358 | 0.339 |
| 4.2 | 0.185 | 0.319 | 0.369 | 0.364 |
| 4.4 | 0.190 | 0.315 | 0.330 | 0.267 |
| 4.6 | 0.185 | 0.319 | 0.330 | 0.305 |
| 4.8 | 0.174 | 0.304 | 0.314 | 0.251 |
| 5 | 0.171 | 0.290 | 0.309 | 0.299 |

\*highest (best-performing) value

### List of Supplemental Files

Supplemental_File_1.zip. Related to Figure 4 and Figure S3. Cytoscape sessions with NetNC-FTI results for TF_ALL.

Supplemental_File_2.zip. Related to Figure 6. Primers for RT-qPCR

